# Response nonlinearities in networks of spiking neurons

**DOI:** 10.1101/856831

**Authors:** Alessandro Sanzeni, Mark H. Histed, Nicolas Brunel

## Abstract

Combining information from multiple sources is a fundamental operation performed by networks of neurons in the brain, whose general principles are still largely unknown. Experimental evidence suggests that combination of inputs in cortex relies on nonlinear summation. Such nonlinearities are thought to be fundamental to perform complex computations. However, these non-linearities contradict the balanced-state model, one of the most popular models of cortical dynamics, which predicts networks have a linear response. This linearity is obtained in the limit of very large recurrent coupling strength. We investigate the stationary response of networks of spiking neurons as a function of coupling strength. We show that, while a linear transfer function emerges at strong coupling, nonlinearities are prominent at finite coupling, both at response onset and close to saturation. We derive a general framework to classify nonlinear responses in these networks and discuss which of them can be captured by rate models. This framework could help to understand the observed diversity of non-linearities observed in cortical networks.

**AUTHOR SUMMARY:** Models of cortical networks are often studied in the strong coupling limit, where the so-called balanced state emerges. In this strong coupling limit, networks exhibit without fine tuning, a number of ubiquitous properties of cortex, such as the irregular nature of neuronal firing. However, it fails to account for nonlinear summation of inputs, since the strong coupling limit leads to a linear network transfer function. We show that, in networks of spiking neurons, nonlinearities at response-onset and saturation emerge at finite coupling. Critically, for realistic parameter values, both types of nonlinearities are observed at experimentally observed rates. Thus, we propose that these models could explain experimentally observed nonlinearities.

## INTRODUCTION

The ability of the brain to perform complex functions relies on circuits combining inputs from several sources to make decisions and drive behavior. The principles governing how different inputs are combined in neuronal circuits have yet to be uncovered. One of the leading theoretical models for cortical dynamics and its dependence on inputs is the balanced network model [1; 2]. In this model, a ‘balanced state’ in which excitation and inhibition approximately cancel each other emerges dynamically, without fine tuning, in the strong coupling limit. This model captures in a parsimonious fashion multiple aspects of cortex dynamics. In particular, it leads to irregular firing [1; 2; 3; 4], wide firing rate distributions [1; 2; 3; 5], and weak correlations [6] (but see [7; 8; 9]). These properties match experimental observations [6; 10; 11; 12; 13; 14; 15] and seem to be universal features of strongly coupled units, as they emerge in strongly coupled networks of binary units [1; 2], current-based spiking neurons [4; 5], and conductance-based spiking neurons [16]; although the underlying operation mechanism might differ [16]. Another feature of strongly coupled neural networks is their linear input output relationship, or network transfer function [1; 2; 4; 16]. This property, however, is problematic for different reasons. First, it limits the possible computations implementable by such networks, as nonlinearities are fundamental to perform computations, and layers of networks with linear transfer functions can only perform linear computations. Second, it implies that different inputs should be summed linearly; a prediction which is contradicted by experimental evidence in cortex. In fact, multiple studies have found that neural responses to preferred stimuli are suppressed by contextual stimuli at high contrast and enhanced at low contrast (e.g. [17; 18; 19; 20]). Third, linear combination of inputs fails to predict responses to natural images starting from those measured with elementary stimuli [21].

Different mechanisms have been proposed to explain how nonlinearities can be produced in networks of neurons. One possibility is that nonlinear network response are generated by short-term plasticity [22]. However, the degree to which synapses are facilitated or depressed in vivo is not known. Moreover, it would be informative to understand whether nonlinear computations can be produced in networks with linear synapses. It has been pointed out that nonlinearities can be produced in rate models featuring a power-law transfer function [23; 24]. In particular, it has been shown that these models can produce saturated response while preserving contrast invariant tuning [23], and that these features are reproducible in simulations of conductance-based neurons [23]. In the stabilized supralinear network model (or SSN [24]), power-law transfer functions have been used to explain a variety of nonlinearities observed in cortex [25]. While these papers have been successful in explaining experimental data, their relationship to more realistic networks of spiking neurons is not fully understood. More generally, a theory of nonlinearities in networks of spiking neurons, explaining under what conditions they can be generated and what are the underlying mechanisms, is missing.

In this paper, we investigate analytically and numerically the response of networks of current-based integrate-and-fire neurons as a function of coupling strength. We show that, while a linear transfer function is obtained in the strong coupling limit, nonlinearities at response-onset and at saturation appear as the coupling is decreased. We systematically characterize how they are shaped by single neuron properties and network connectivity, and compare them to nonlinearities observed in rate models with power-law transfer function. A preliminary version of these results has been presented as abstract in [26].

## MATERIALS AND METHODS

### Networks of spiking neurons

We study a randomly connected network of excitatory and inhibitory leaky integrate-and-fire neurons, using a theoretical framework that was developed in [3] and [4]. The network is composed of *N* leaky integrate-and-fire (LIF) neurons, out of which *N*_*E*_ are excitatory (E) and the remaining *N*_*I*_ = *N* − *N*_*E*_ are inhibitory (I). The dynamics of the membrane potential of neuron *i* (*i* = 1, …, *N*) obeys

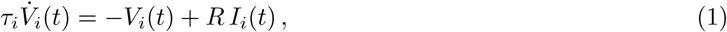

where *τ*_*i*_, *R* and *I*_*i*_ are the membrane time constant, the membrane resistance and the input current of the neuron. The input current is generated by the sum of incoming spikes generated by pre-synaptic neurons, which could be within or outside the network (external inputs); this input is written as

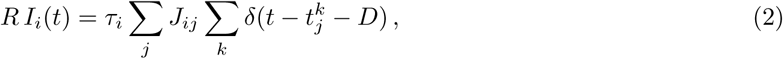

where 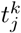 is the *k*-th spike generated by pre-synaptic neuron *j* at time *t* − *D, D* is a synaptic delay, and *J*_*ij*_ is the synaptic efficacy from neuron *j* to neuron *i*. Every time the membrane potential *V*_*i*_ reaches the firing threshold *θ*, neuron *i* emits a spike, its membrane potential is set to a reset *V*_*r*_, and stays at that value for a refractory period *τ*_*rp*_; after this time the dynamics continues as before. Assuming that connectivity is sparse, that a neuron receives a large fixed number of presynaptic inputs from each presynaptic population, that each presynaptic spike causes a small change in membrane potential (*J*_*ij*_*/θ* ≪ 1), that temporal correlations in synaptic inputs can be neglected, that all neurons in a given population are described by the same single cell parameters, and that the network is in an asynchronous state in which all neurons fire at a constant firing rate, the firing rate of neurons in population *A* (*A* = *E, I*) is given by [4]

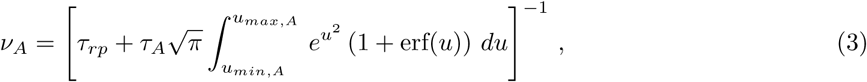

where *u*_*max,A*_ and *u*_*min,A*_ are the distance form threshold and reset of the mean input *μ*_*A*_ measured in units of noise *σ*_*A*_, i.e.

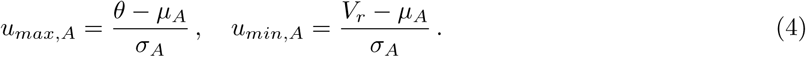

where means and variances are given by

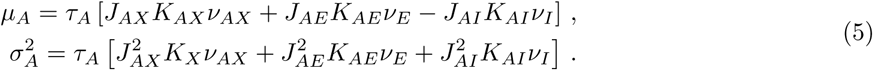

Here *ν*_*AX*_ are the average firing rates of neurons from outside the network providing inputs to population *A*, while *K*_*AB*_ and *J*_*AB*_ are the number of connections and the synaptic efficacy from population *B* (*B* = *E, I* and *X* for external) to population *A*. The right side of Eq. (3) is sometimes referred to as the Ricciardi transfer function, or nonlinearity [27]. It relates the presynaptic input mean *μ* and noise *σ* of a neuron to its firing rate.

Firing variability is quantified with the coefficient of variation (CV) of inter-spike intervals (ISI), given by [4]

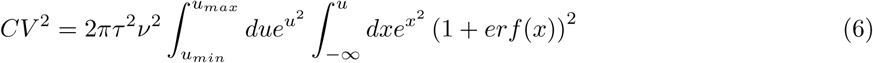

The network activity is found self-consistently from Eqs. ((3)-(5). As in [4] we investigated two models: model A, in which all neurons have the same biophysical and input connectivity properties, and model B, in which the excitatory and the inhibitory populations have different properties.

Through the paper, we use the following parametrizations:

- In model A, we use *J*_*EE*_ = *J*_*IE*_ = *J, J*_*EX*_ = *J*_*IX*_ = *g*_*X*_*J, J*_*EI*_ = *J*_*II*_ = *gJ, K*_*EX*_ = *K*_*EE*_ = *K*_*IX*_ = *K*_*IE*_ = *K* and *K*_*EI*_ = *K*_*II*_ = *γK*, so that the equations for means and variances become (note that in this model the excitatory and inhibitory rates are equal, *ν*_*E*_ = *ν*_*I*_ = *ν*)

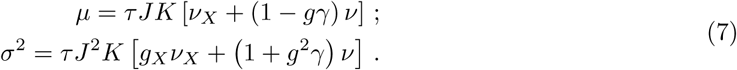

where, without loss of generality, we have absorbed a factor *g*_*X*_ in the external drive *ν*_*X*_.
- In model B, we use *J*_*EX*_ = *g*_*EX*_*J*_*EE*_, *J*_*IX*_ = *g*_*IX*_*J*_*IE*_, *J*_*EI*_ = *g*_*E*_*J*_*EE*_, *J*_*II*_ = *g*_*I*_*J*_*IE*_, *K*_*EX*_ = *K*_*EE*_, *K*_*IX*_ = *K*_*IE*_ and *K*_*AI*_ = *γK*_*AE*_. In this case, the equations for means and variances in both populations (*A* = *E, I*) read

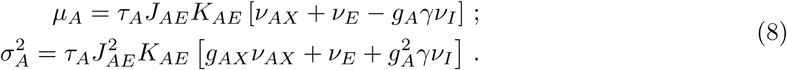

where we have again absorbed *g*_*AX*_ in the external drive *ν*_*AX*_. Moreover, we take inputs of the form *ν*_*AX*_ = *α*_*A*_*ν*_*X*_, where *α*_*E,I*_ are fixed parameters while *ν*_*X*_ represents the intensity of the input.

Throughout the paper, we use *θ* = 20mV, *V*_*r*_ = 10mV, *τ*_*E*_ = *τ*_*I*_ = 20ms, *τ*_*rp*_ = 2ms, and *γ* = 0.25. Note that the equations for the firing rates are independent on the synaptic delays *D* and the total number of neurons *N*. We specify the values of these parameters for Fig. 8 where our analytical results are compared with numerical simulations.

**FIG. 1.**
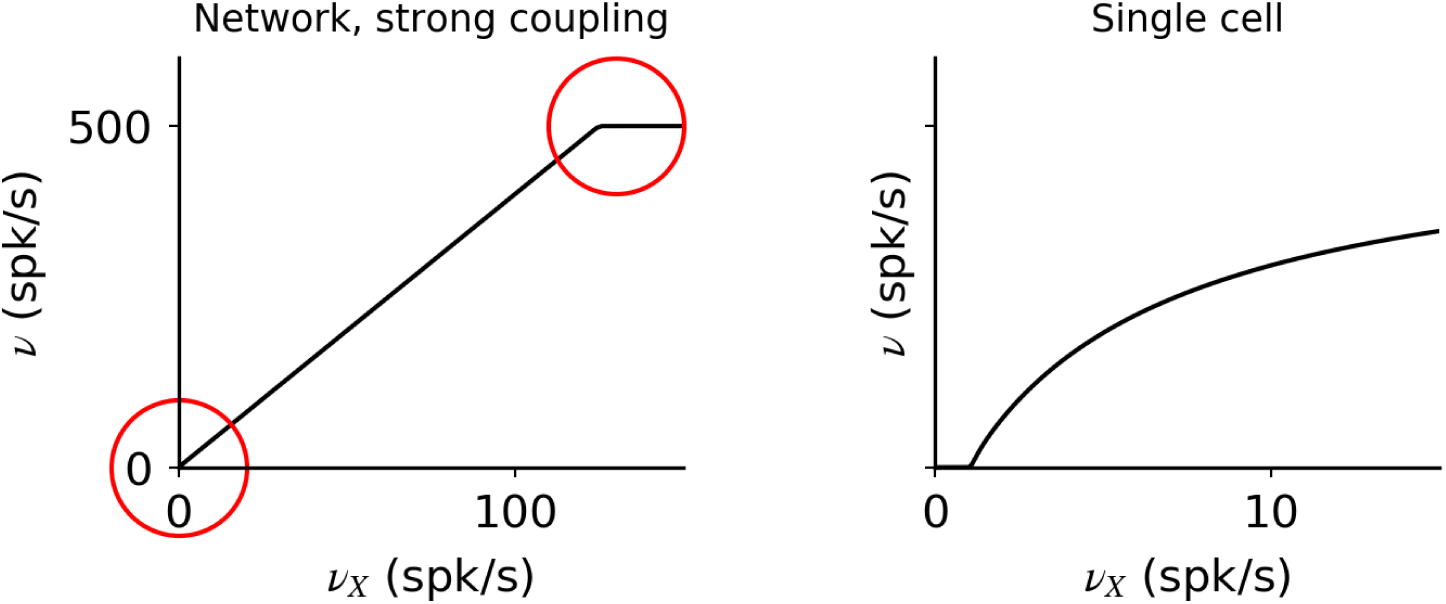
Network transfer function in the strong and weak coupling limits. Network response in strongly coupled networks (left) features linear regions separated by abrupt transitions (red circles). In the weak coupling limit, the network transfer function becomes identical to the single cell transfer function. It is supralinear at low inputs, but then becomes sublinear at high inputs.

**FIG. 2.**
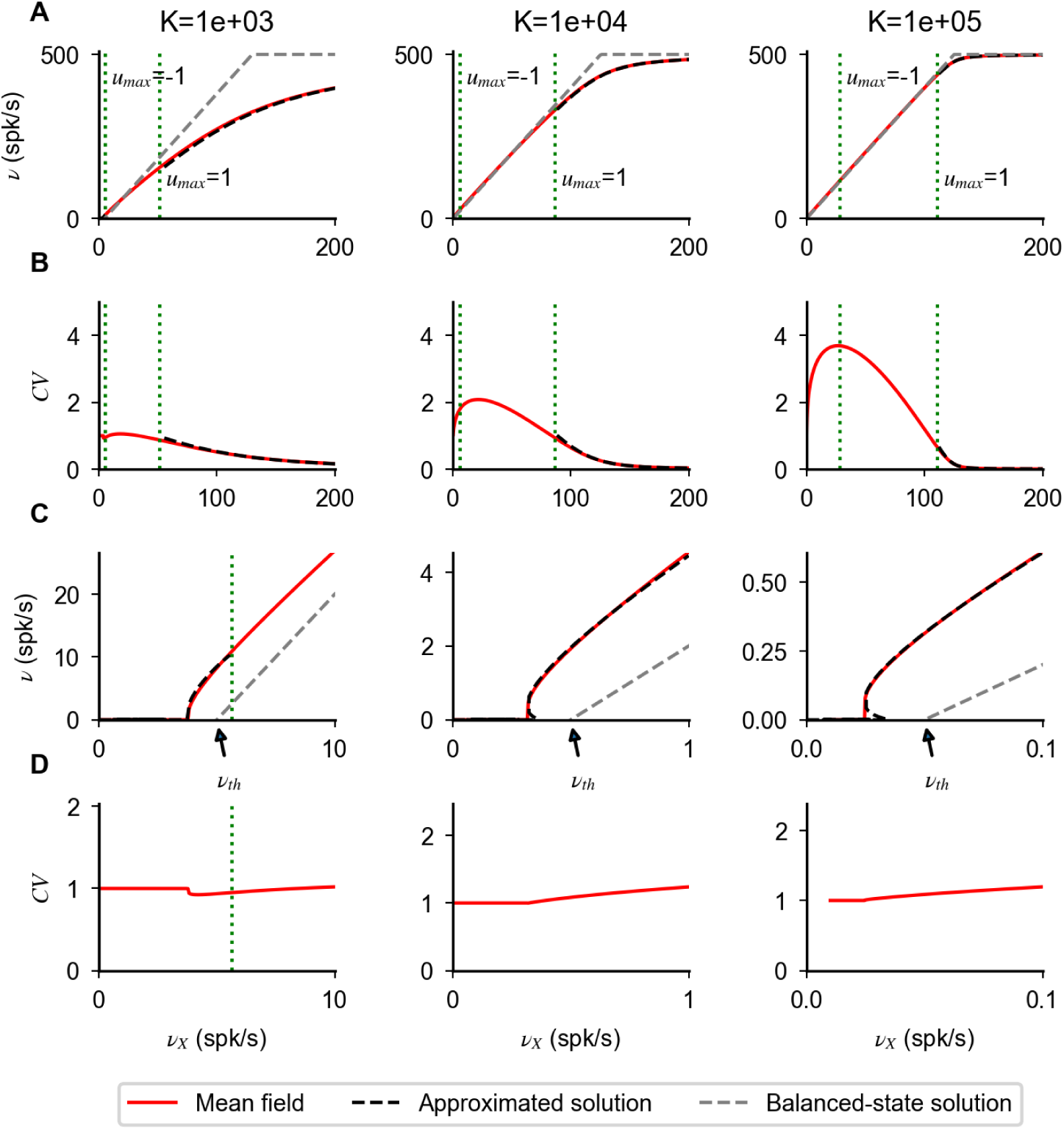
Types of nonlinearities in network response. (**A**) Network transfer function obtained solving numerically Eqs. (3)-(5) for different values of *K* (indicated on top of each column). (**B**) CV of interspike interval distribution obtained solving numerically Eq. (6) (red lines). (**C**,**D**) Plots as in **A, B** but zoomed in the region of response onset. In all panels, dotted green lines correspond to the values at which *u*_*max*_ = 1 and *u*_*max*_ = −1, i.e. they indicate the separation between the different operating regimes mentioned in the main text. Firing nonlinearities at response onset and saturation are captured by approximated forms (black dashed lines) obtained for *u*_*max*_ ≪ −1 (Eq. (15)) and *u*_*max*_ ≫ 1 (Eq. (13)), respectively. For *u*_*max*_ ∼ 0, the transfer function approaches the balanced-state solution (gray dashed lines, Eq. (17)); in the region −1 < *u*_*max*_ < 1, the first order corrections are of order 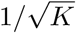 and become negligible as *K* increases. In the suprathreshold regime (*u*_*max*_ ≪ −1), the *CV* approaches zero, i.e. firing becomes regular, with a decay with input strength captured by Eq. 14 (dashed line). Parameters: *J* =0.2 mV, *g*=5.0, *g*_*X*_ =1.

**FIG. 3.**
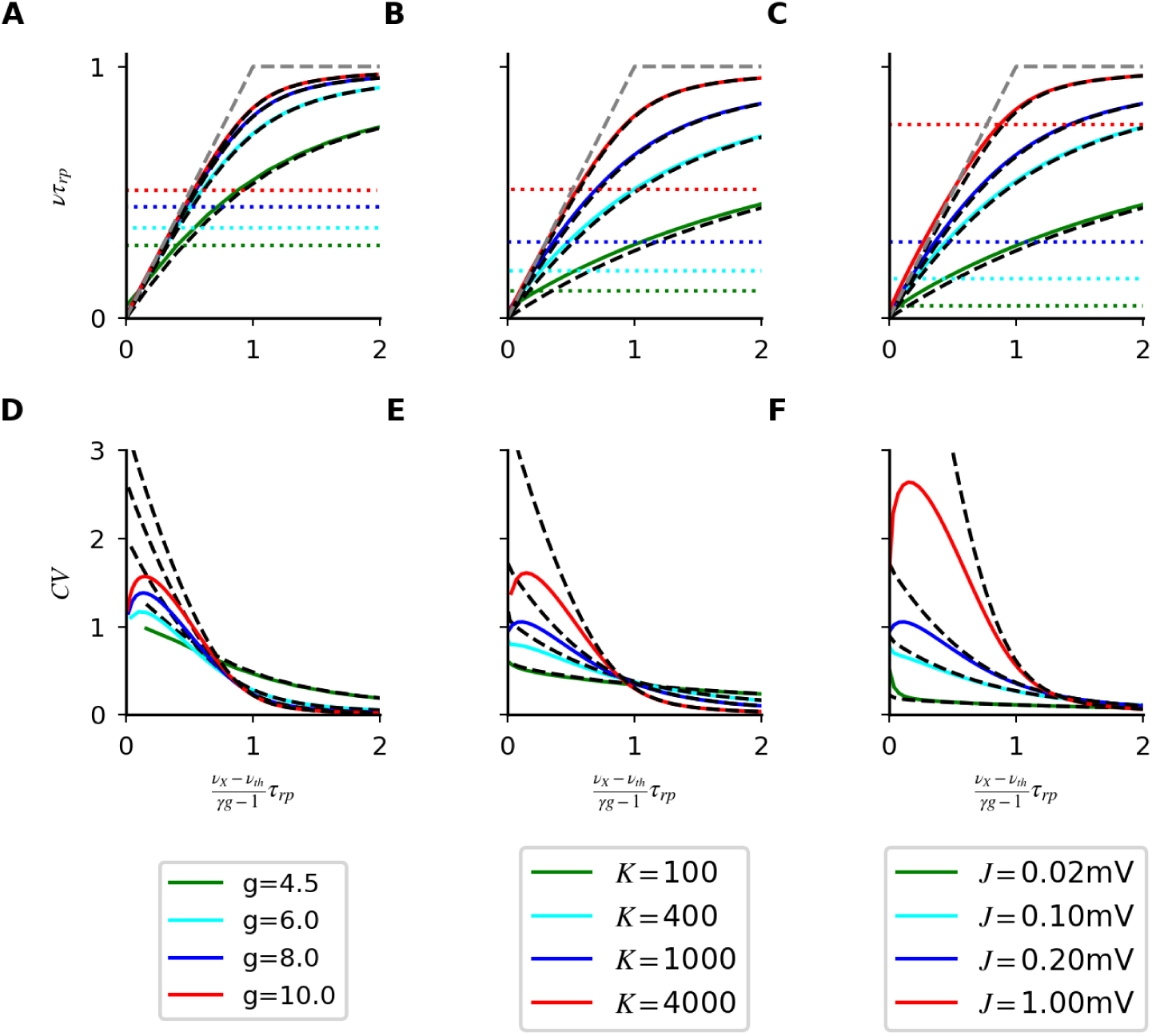
Saturation nonlinearities in Model A. Transfer function (first row) and *CV* (second row) computed numerically from Eq. ((3)-(5)) and Eq. (6) (continuous lines) for different *g* (first column), *C* (second column) and *J* (third column). As in Fig. 2, colored dotted lines in the first row represent values of the rates at which *u*_*max*_ = −1. Black dashed lines solutions of the approximated Eqs. (12) and (14). This validation of the approximated rate equation motivates the perturbative approach used in the main text and allows to classify nonlinearities in a general way. Simulation parameters: *J* =0.2 mV, *K* = 10^3^ in (**A**,**D**); *g*=5.0, *J* = 0.1mV in (**B**,**E**); *g*=5.0, *K* = 10^3^ in (**C**,**F**); in all plots *g*_*X*_ =1, *τ* =20ms.

**FIG. 4.**
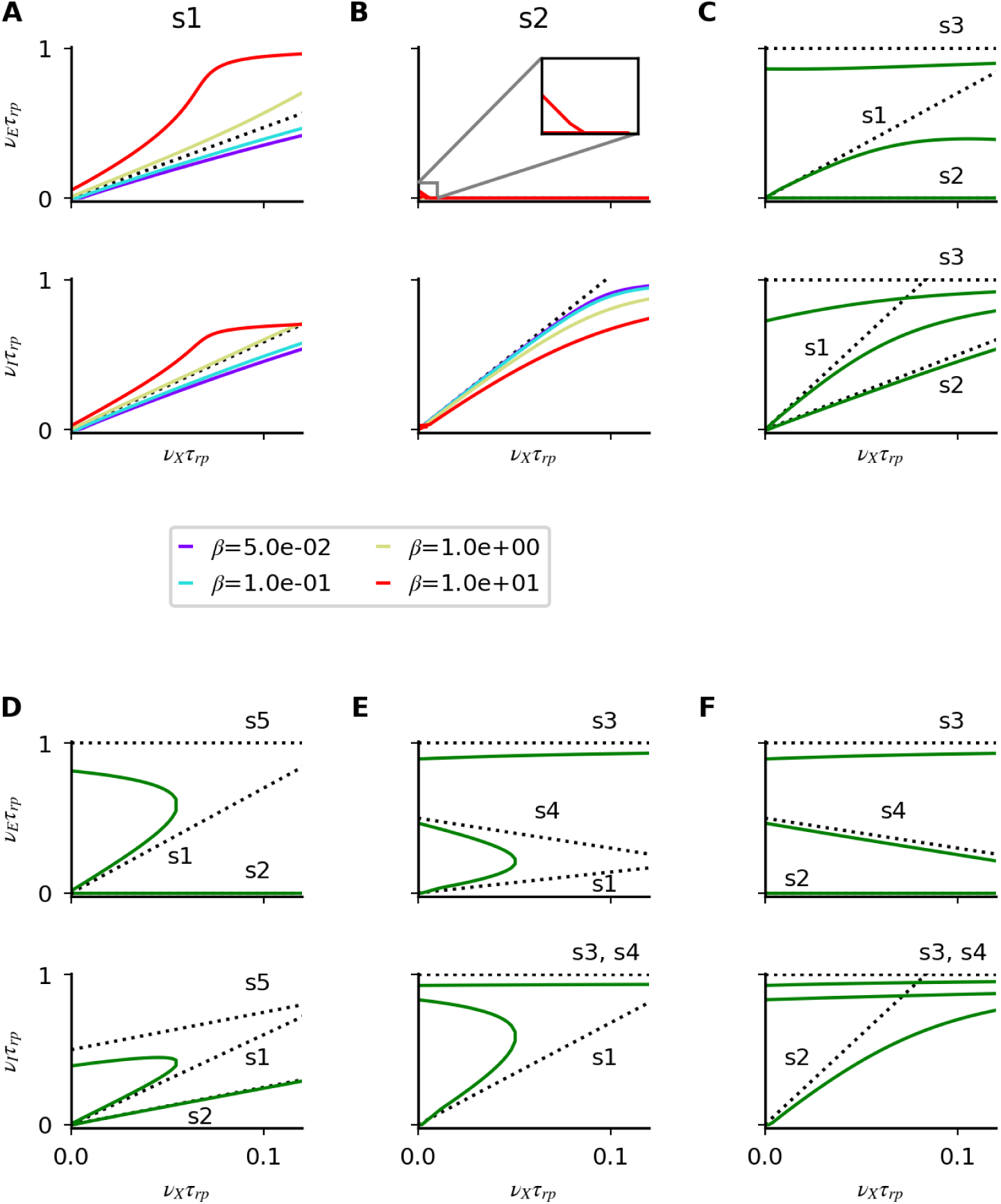
Saturation nonlinearities in model B. (**A**-**F**) Numerical solutions of Eq. (3)-(5) for different parameters (colored lines) and linear approximations predicted by Eqs. (26) and (27) in the strong coupling limit (dotted lines). In each panel, the first (second) row shows the excitatory (inhibitory) firing rate as a function of *ν*_*X*_. (**A**,**B**) Nonlinear solutions obtained at finite coupling starting from solution s1 and s2 for different *β* values. (**C**-**F**) All admissible cases of coexistence of multiple solutions at low *ν*_*X*_; note that, as expected from our analysis, the number of solution changes as *ν*_*X*_ increases. Simulation parameters are: (**A**) *g*_*I*_=3.9, *g*_*E*_=8, *α*_*I*_=1 *α*_*E*_=7; (**B**) *g*_*I*_=3.9, *g*_*E*_=8, *α*_*I*_=10, *α*_*E*_=7; (**C**) *g*_*I*_=4, *g*_*E*_=3, *α*_*I*_=5, *α*_*E*_=2; (**D**) *g*_*I*_=8, *g*_*E*_=6, *α*_*I*_=5, *α*_*E*_=2; (**E**) *g*_*I*_=1, *g*_*E*_=2, *α*_*I*_=0.3, *α*_*E*_=2; (**F**) *g*_*I*_=1, *g*_*E*_=2, *α*_*I*_=3, *α*_*E*_=2. In panels (**A**-**F**), *g*_*EX*_ = *g*_*IX*_ = 1; *J*_*EE*_ = *J*_*IE*_ = 0.2*mV* and *K*_*EE*_ = *K*_*IE*_ = 10^3^; except in **A** and **B** where 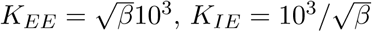.

**FIG. 5.**
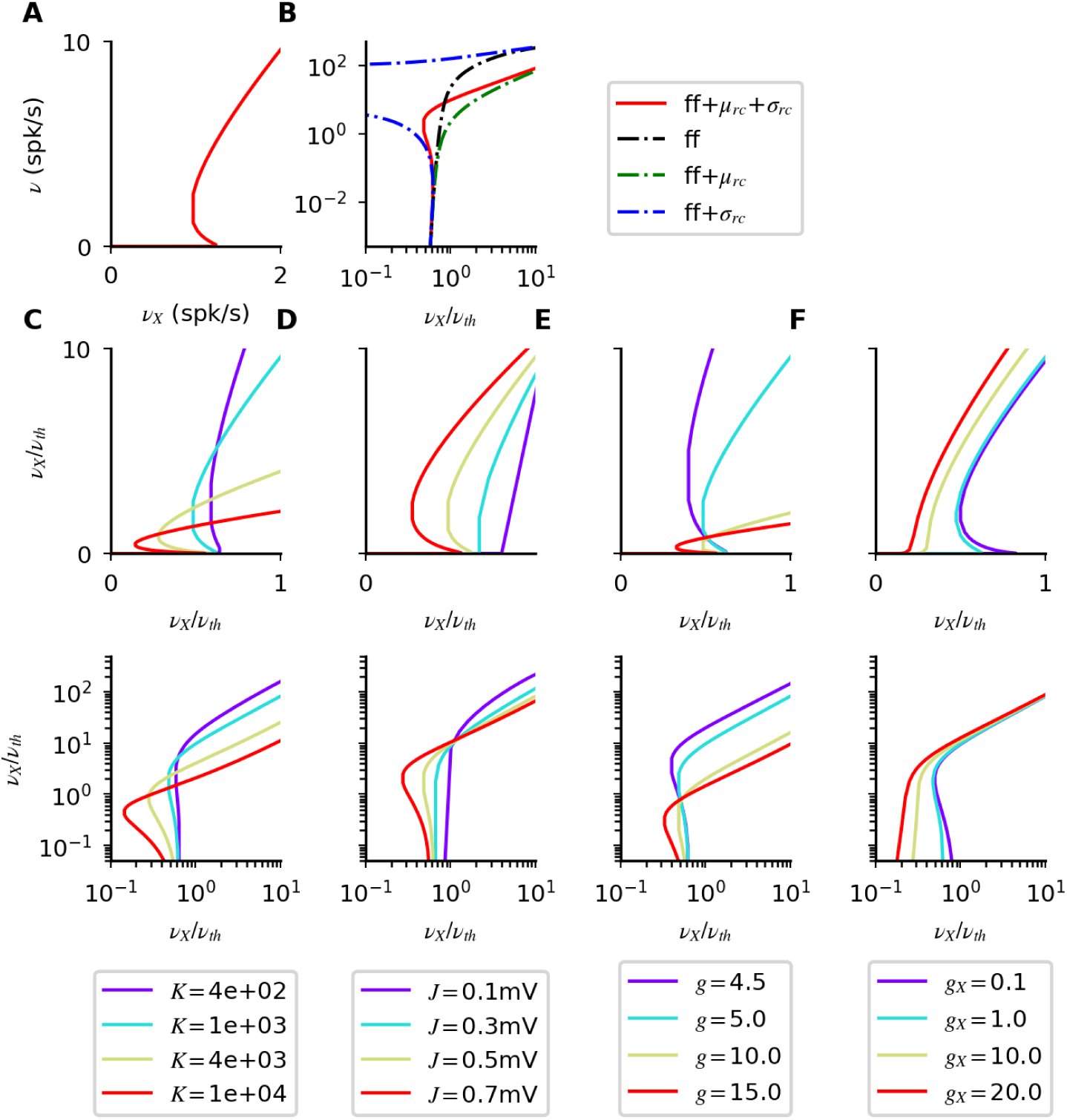
Response-onset nonlinearities in model A. (**A**-**B**) Network response computed from Eq. ((3)-(5)) (red). Response at low rates is determined only by feedforward inputs (black, Eq. (30)); effects of recurrent noise (blue) and mean (green) become relevant at higher rates, with opposite effects on the number of solutions. (**C-F**) Effects of different parameters on the number of solutions in network response Simulation parameters, unless otherwise specified in legend, are *K* = 10^3^, *J* =0.5 mV, *g*=5.0, *g*_*X*_ =1, *τ* =20ms.

**FIG. 6.**
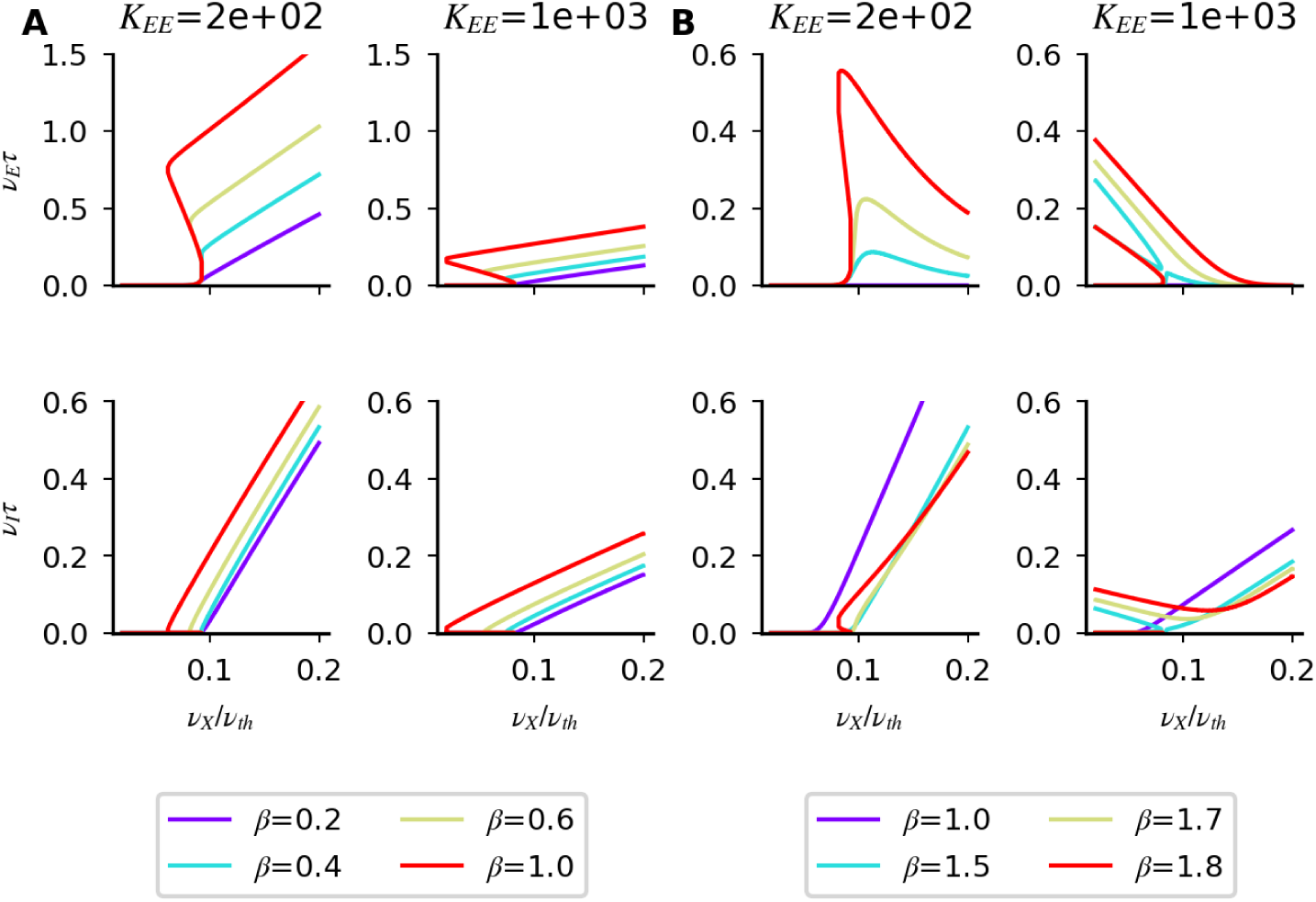
Onset-nonlinearities in model B. Responses obtained solving numerically Eq. (3)-(5) for regular (**A**) and supersaturated (**B**) solutions at different coupling strengths and for different *β* = *K*_*EE*_*J*_*EE*_*/K*_*IE*_*J*_*IE*_. The size of the nonlinear region decreases as the coupling strength increases and as the inhibitory activation threshold decreases. In B, the lack of purple curve in the first row means that the firing rates are very close to zero in the whole range of inputs. In the plots, response onset emerges around 0.1 *ν*_*th*_ because of the value *g*_*EX*_ = *g*_*IX*_ = 10 used in simulations. As showed in Fig. 5F, this choice enhances the amplitude of the nonlinear region by decreasing the likelihood of having multiple solutions. Other simulation parameters are:, *J*_*EE*_ = *J*_*IE*_ = 0.1mV, *g*_*I*_=3.9, *g*_*E*_=8. In (**A**) *α*_*I*_=1 *α*_*E*_=7; in (**B**) *α*_*I*_=10, *α*_*E*_=7.

**FIG. 7.**
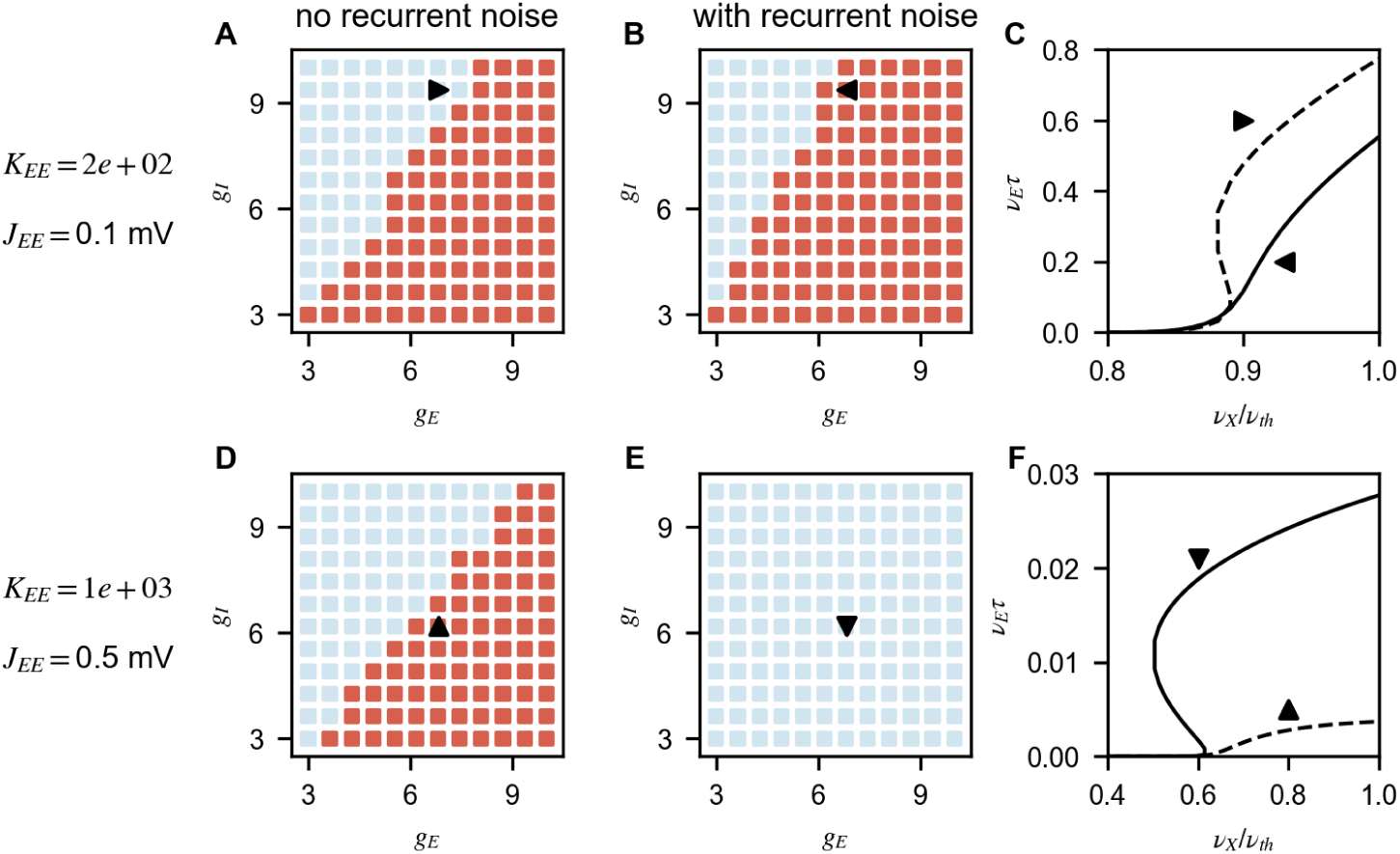
Role of recurrent noise in generating multiple solutions at onset in model B. Number of solutions for different combinations of *g*_*E*_ and *g*_*I*_ (red and blue correspond to single solution and multiple solutions, respectively) computed without (first column) and with (second column) recurrent noise; the third column shows transfer functions for specific combinations (indicated by triangles in the first two columns). Both for small (first row) and large (second row) noise influences the number of solution: (**A**-**C**) For some parameters, noise reduces the size of the region with multiple solutions. (**D**-**F**) For others, noise increases the size of the region with multiple solutions. Parameters: *J*_*IE*_ = *J*_*EE*_, *g*_*IX*_ = *g*_*EX*_ =1, *α*_*I*_ = *α*_*E*_ = 1.

**FIG. 8.**
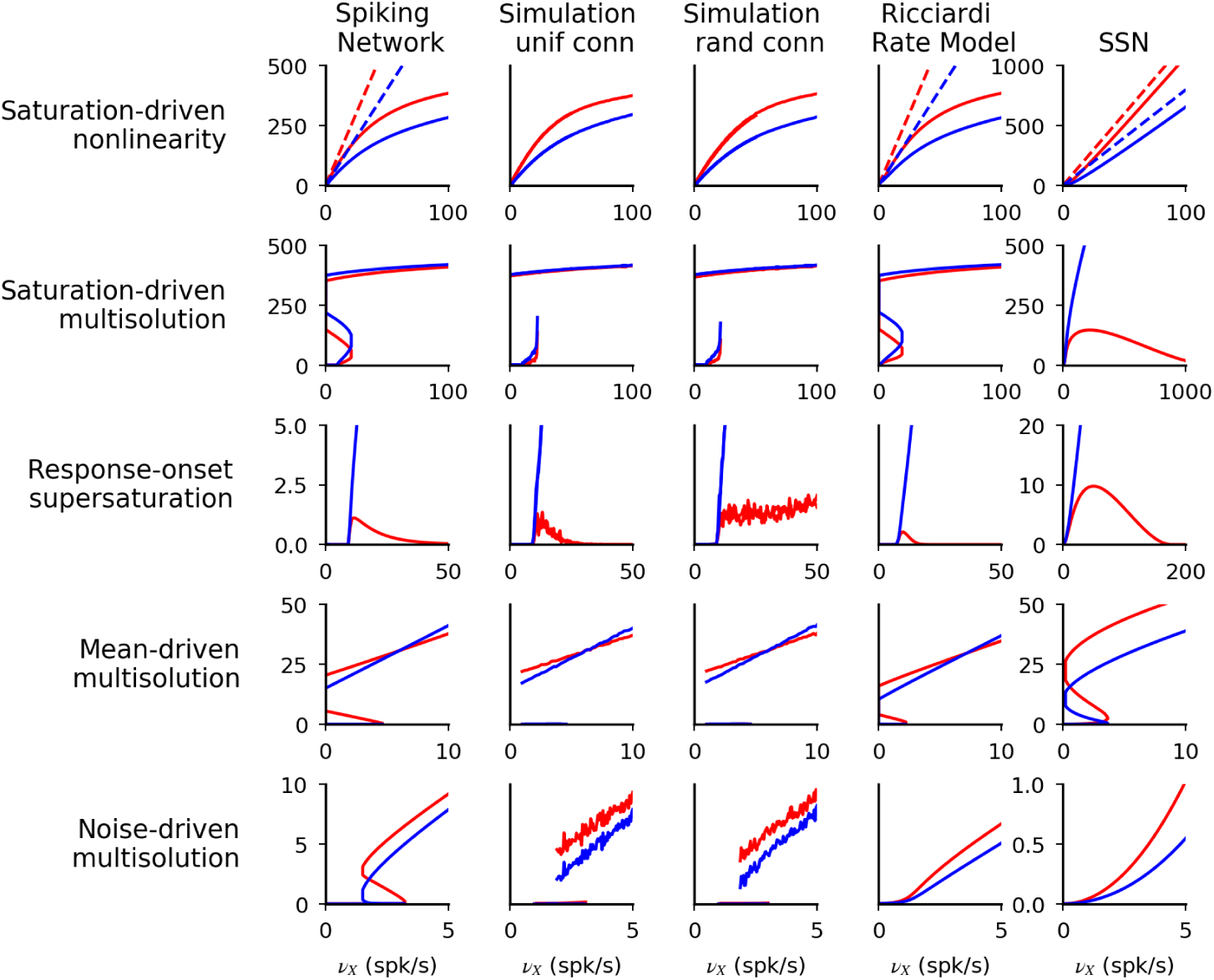
Spiking-networks nonlinearities in network simulations and rate models. Comparison of responses computed with: mean field theory of spiking networks (first column); network simulation with uniform (second column) and random (third column) connectivity; rate models (Ricciardi model, fourth column, and the SSN, fifth column). Different rows correspond to different values of *α*_*E,I*_ and *g*_*E,I*_. Prediction of the mean field theory match numerical simulations, with the only exception of the lack of supersaturation in networks with random connectivity. Spiking networks present nonlinearities at saturation and at response-onset. Saturation nonlinearities are generated by the refractory period and hence are captured only by the Ricciardi model and not by the SSN, which lack this ingredient. These nonlinearities have effects also at low rate and constraints the parameter space over which the response is unique. Response-onset nonlinearities have similar structure in the three models but in spiking network are smaller in size and feature multiple solutions generated by noise feedback. Network structure (from top): *g*_*E*_ = 8, *g*_*I*_ = 7, *α*_*E*_ = 4, *α*_*I*_ = 2; *g*_*E*_ = 2.08, *g*_*I*_ = 1.67, *α*_*E*_ = *α*_*I*_ = 1; *g*_*E*_ = 4.5, *g*_*I*_ = 2.9, *α*_*E*_ = *α*_*I*_ = 1; *g*_*E*_ = 4.1, *g*_*I*_ = 2.46, *α*_*E*_ = 1, *α*_*I*_ = 0.2; *g*_*E*_ = 7, *g*_*I*_ = 6, *α*_*E*_ = 1, *α*_*I*_ = 0.7. In all simulations, for spiking-networks mean field, simulations and Ricciardi model: *J*_*EE*_ = *J*_*IE*_ = *J*; *K*_*EX*_ = *K*_*EE*_ = *K*_*IE*_ = *K*; *g*_*EX*_ = *g*_*IX*_ = 1. (except second row, where *J*_*EE*_ = *J*_*IE*_ 2.5/2.4 = *J*, and fourth row, where *K*_*EX*_ = *K*_*EE*_ = 2*K*_*IE*_ = 2*K*); values are for 400 *K* and 0.2mV for *J* in all simulations except for noise driven bistability where *J* = 0.5mV. In Ricciardi model, *σ*_*E*_ = *σ*_*I*_ = *σ*; *σ* matches noise at threshold in spiking neurons and is (from the top): 25mV, 10mV, 3mV, 5mV, 7mV. In the SSN, *k* = 0.04, *n* = 2, *W*_*EE*_ = *W*_*IE*_ = 1, *W*_*AI*_ = *γg*_*A*_ except in second row, where *W*_*IE*_ = 2.4/2.5 and *W*_*EI*_ = *γg*_*I*_ 2.4/2.5, and fourth row where *W*_*EE*_ = 2 and *W*_*EI*_ = 2*γg*_*E*_.

### Rate models

In the last part of the results section, we analyzed which of the nonlinearities observed in spiking networks emerge in rate models. We study this question using two specific rate models:

#### Ricciardi rate model

: It is defined here using the single neuron transfer function of Eq. (3), assuming the noise amplitude is fixed. The *f* − *μ* curve of the *A* population is then defined as

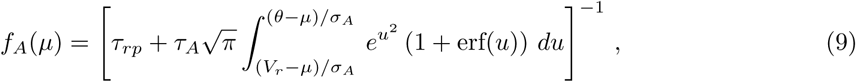

where *σ*_*A*_ is a fixed parameters. Once the *f* − *μ* curve has been fixed, rates in the network are computed as above, i.e. a self-consistency condition is imposed which gives the mean input currents *μ*_*A*_s as a function of the external drive and recurrent interactions.

#### Supralinear Stabilized Network (SSN) rate model

: In the SSN model, the *f* − *μ* curve is given by a power-law

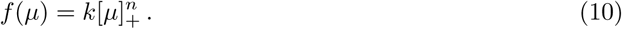

Rates are computed as for the Ricciardi rate model.

## RESULTS

### Two nonlinear regions in network response at finite coupling

The main goal of this paper is to characterize systematically the network transfer function, that describes how the average E and I firing rates depend on external inputs to the network, and in particular how this transfer function depends on network parameters.

This transfer function is well understood in two opposite limits (see Fig. 1 for an illustration of these two limits):

1. In the strong coupling limit, excitatory and inhibitory mean inputs need to ‘balance’ in both E and I neurons, for total inputs to remain finite, leading to a linear relationship between output and input rates [1; 2]. This linear relationship can only hold provided specific conditions on network parameters are satisfied (see below). Furthermore, this linear relationship only holds in a limited range of inputs, since output firing rates must be bounded between zero and a maximal value imposed by the refractory period. Therefore, the network transfer function is piecewise linear, as shown in Fig. 1A.
2. In the opposite weak coupling limit, the recurrent inputs become negligible compared to external inputs. Therefore, the network transfer function becomes in this limit identical to the single neuron transfer function, Eq. (3). In the presence of fluctuations in external inputs, this transfer function is expected to be generically sigmoidal (see Fig. 1B), with a supralinear region at low rates, in the so-called subthreshold, or fluctuation-driven regime, while it is expected to become sublinear at higher rates, because of the refractory period (e.g. [28]).

Intuitively, we therefore expect that recurrent connections should lead to an interpolation between these two extremes. In particular, we expect two non-linear regions, one at low rates, one at high rates (shown as red circles in Fig.1A), whose size should decrease as coupling strength increases, separated by a linear region, whose size should increase as coupling strength increases. Furthermore, we would expect naively that the low rate region is supralinear, while the high rate region is sublinear. In the following, we show that these naive expectations are not necessarily true, since other behaviors are possible.

To get more insight into what controls reponse non-linearities, we need to analyze the equations that give the average population firing rates as a function of network parameters, Eqs. (3-5). Here, we start by considering model A, in which E and I neurons have identical parameters, leading to a single equation describing the firing rates of both populations. The solutions of this equation depend on the single neuron transfer function, Eq. (3). In particular, Eq. (3) tells us that the mean firing rate is given as a function of two variables, *u*_*max*_ = (*θ* − *μ*)*/σ* and *u*_*min*_ = (*V*_*r*_ − *μ*)*/σ*, that describe the distance of the mean inputs to neurons from threshold and reset, respectively, in units of input noise *σ*.

#### The high input/high rate regime

When the mean inputs are far above threshold (*u*_*max*_ ≪ −1), neural firing is dominated by deterministic drift in the neuron membrane potential, and noise has only a weak effect on firing. In this regime, we use the first order term in the expansion

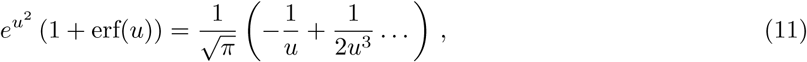

valid for large and negative *u*, to obtain a simplified expression of Eq. (3),

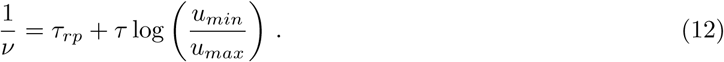

Eq. (12), is the transfer function of a single neuron receiving deterministic input, as expected from the fact that the mean input is far above threshold in units of the input noise. Higher order terms in expansion (11) provide corrections due to membrane fluctuations. Eq. (12) shows that the relation between *ν* and *ν*_*X*_ in this regime is nonlinear. With the additional assumption |(*u*_*max*_ − *u*_*min*_)*/u*_*max*_| ≪ 1 (which is equivalent to *θ* − *V*_*r*_ ≪ *μ* − *θ*, a condition satisfied for *μ* sufficiently above threshold *θ*), we can expand Eq. (12) and obtain a direct relation between *ν* and *ν*_*X*_

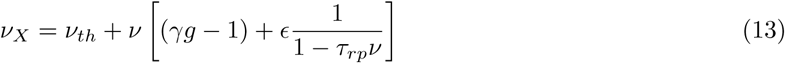

where 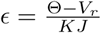 is the fraction of recurrent excitatory inputs that are needed to fire simultaneously to drive the membrane from reset to threshold, and *ν*_*th*_ = *θ/KJτ* is the external input needed to generate spikes in the absence of noise, respectively. Eq. (13) shows explicitly that the nonlinearity in this regime is generated by the presence of the refractory period *τ*_*rp*_, while the parameter *E* controls the deviation from the linear prediction.

In Fig. 2 first row, we compare the prediction of Eq. (13) with the numerically computed solutions of Eqs. (3)-(5) for various values of *K* = 1000, 10, 000, 100, 000, *J* = 0.2mV, *g* = 5. Numerical solutions of the mean-field equations show that our approximation gives a good description of the transfer function, capturing the nonlinearity observed close to saturation. As expected from Eq. (13), the width of the nonlinear region close to saturation expands as *E* increases. For instance, for *K* = 1, 000, we have *ϵ* = 0.05, and the deviations from linear behavior are already significant when the firing rate is less than half its maximal value. On the other extreme, for the unrealistically high value *K* = 100, 000, *ϵ* = 0.0005 and the non-linear region becomes extremely small.

Consistent with the idea that, for *u*_*max*_ ≪ −1, firing is driven by deterministic drift, the CV of the ISI is found to be smaller than one in the high input regime (see Fig. 2 second row); with a value that becomes smaller and smaller as *ν*_*X*_ increases. Using Eq. (11), we derive a simplified expression for the CV starting from Eq. (6) given by

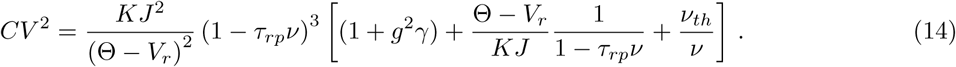

This expression captures the decay of the CV observed in Fig. 2, and shows that, as the neural firing rate *ν* approaches its maximum value, the *CV* goes to zero as 1 − *τ*_*rp*_*ν*.

#### Low input/low rate regime

For membrane potential far below threshold (*u*_*max*_ ≫ 1), firing is driven by large stochastic fluctuations, so that noise can no longer be neglected. In this regime, in the simplified framework of model A, Eq. (3) is well approximated by [4]

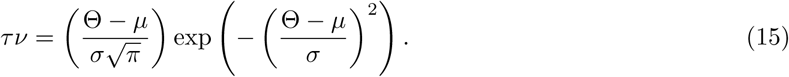

The transfer function obtained solving Eq. (15) is nonlinear and provides a good description of the response for small rates. Specifically, the response shows threshold like nonlinearities, with threshold close to *ν*_*th*_; we will characterize this in more detail in the following sections. Consistent with the fact that firing is produced by large fluctuations, the response is highly irregular (CV of order one or above) (see Fig. 2 second and fourth row). Note that the CV depends non-monotonically on input rate, as shown in [29].

#### Intermediate linear region

Up to now, we have shown that network response is expected to be nonlinear in the regions of rates corresponding to *u*_*max*_ ≪ −1 and *u*_*max*_ ≫ 1; we will now show that in the region between these two regimes the response is expected to be approximately linear. For fixed values of *ν*_*X*_ and *ν*, we can write *u*_*max*_ = *ω* with *ω* constant of order one. In this regime, neural firing is driven by input fluctuations if *ω* > 0, or a combination of deterministic drift and fluctuations if *ω* < 0. The transfer function can be found iteratively for every value of *ν* and *ν*_*X*_ as

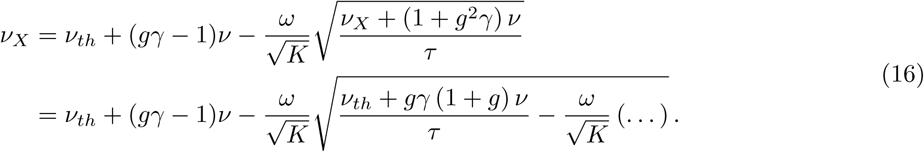

Eq. (16) shows that, for *ω* = 0, the transfer function always matches the balanced-state solution [1; 4]

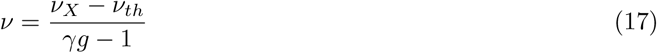

For *ω* of order one, deviation from this solution are expected to be of order 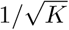. To test this, we plot in Fig. 2 first row the balanced-state solution (gray dashed-lines): network responses are found to be close to this solution, with a distance that decreases as *K* increases. The balanced-state model [1] is characterized by a linear transfer function (given by Eq. (17) with our notation). Consistent with the fact that this model has been derived in the strong coupling limit, we find that as *K* increases the network response converges to the balanced-state solution.

The above argument can be generalized for the more general case of model B. Using the approximations derived above, we can deduce that nonlinearities in the network response will appear any time that one or both populations have mean inputs far below threshold or above threshold. Moreover, an approximately linear solution appears when both mean inputs are close to threshold, with a scaling close to the one predicted by the balanced-state model up to corrections of order 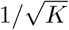. In the following sections we systematically classify these nonlinearities as a function of the network parameters.

### Saturation nonlinearities

In this section, we use a perturbative approach to characterize saturation nonlinearities, first in model A and then in model B. We find that a key role is played by linear solutions obtained in the strong coupling limit, i.e. balanced-solutions, which serves as a starting point for the perturbative expansion, and by the coupling strength, which determines the amplitude of the deviation from linear response.

#### Model A

We start our analysis of saturation nonlinearities by computing numerically responses in the mean field theory, i.e. solving Eq. (3)-(5), for different parameter values. The network response, for mean input *μ* above threshold, depends on *g, K*, and *J*; it depends only weakly on *g*_*X*_ since, in this regime, responses depend weakly on noise amplitude. Results of the numerical analysis are shown in Fig. 3. Note that in this figure input firing rates are rescaled so that the balanced limit is the same regardless of the parameters. The transfer function shows a sublinear scaling as the network rate approaches saturation (Fig. 3 A-C), but nonlinear effects are seen at low rates, especially for small *K*; the *CV* decreases monotonically for sufficiently large inputs (Fig. 3 D-E). To understand the generality of these results, we turn to the approximated expressions of Eqs. (13) and (14).

As shown in Fig. 3 A-C, solutions of Eq. (13) capture the observed nonlinearities across input strength and parameter values. Surprisingly, they provide a good approximation of the network transfer function also for *u*_*max*_ ≈ 1, i.e. beyond the range of validity of Eq. (13). These results suggest that Eq. (13) can be used to understand nonlinearities in the whole range of activity levels above response onset.

To classify possible nonlinear responses, we solve Eq. (13) using a perturbative expansion in *ϵ*,

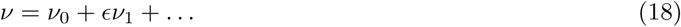

The zeroth order solutions are found taking *ϵ* = 0 (strong coupling limit) in Eq. (13); there are two such solutions:

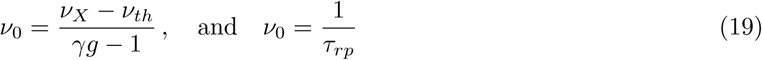

The first solution corresponds to the balanced-state solution of Eq. (17) while the second solution corresponds to saturated activity, with firing at maximum rate.

Using the balanced-state solution in the *ϵ* expansion, we get

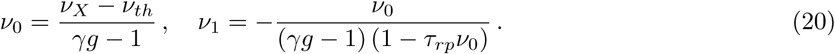

In the inhibition dominated regime (*gγ* > 1), which is thought to underlie cortical dynamics [4], the first order correction is always negative and its absolute value becomes larger as the rate increases; this shows that the output rate increases sublinearly with input rate, regardless of the choice of parameters. Eq. (20) also shows that the deviations from linear response increase with *ϵ*. At finite coupling (*ϵ* > 0), the first order correction is linear if *τ*_*rp*_*ν*_0_ ≪ 1; nonlinear corrections become large when *ϵ* ≈ (*γg* − 1)(1 − *τ*_*rp*_*ν*_0_).

Using the saturation solution in the *ϵ* expansion, we get

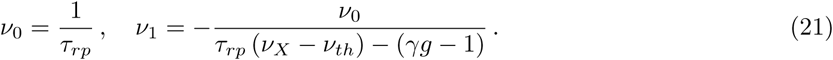

This equation shows that the rate approaches saturation as 1*/ν*_*X*_ for all connectivity parameters. Note that, in the inhibition dominated regime, corrections with *τ*_*rp*_ (*ν*_*X*_ − *ν*_*th*_) < (*γg* − 1) produce rates above saturation, which are not realizable. It follows that the saturation solution appears only when the balanced-solution is larger than 1*/τ*_*rp*_, i.e. the two solutions found at *ϵ* = 0 are mutually exclusive at finite coupling; this will not be true in model B.

As in the previous section, the approximate CV expression of Eq. (14) captures the decay as activity approaches saturation, ensuring that the suppression of irregular firing is expected for all parameter values. However, unlike what happens for the rates, the approximated expression departs from the mean field value as soon as *u*_*max*_ ≈ −1 (see Fig. 3 second row).

#### Model B

In this section, we characterize saturation nonlinearities in model B. Using the perturbative method introduced for model A, we show that the network transfer function, at a fixed external drive, has in some cases multiple solutions. In cases in which there is a unique rate value, depending on the connectivity matrix and coupling strength, the response can be a sub-linear, linear or supra-linear function of the input.

As discussed in the methods section, model B features one excitatory and one inhibitory population with rates *ν*_*E,I*_ and external drive *ν*_*EX*_, *ν*_*IX*_ = *α*_*E,I*_*ν*_*X*_. The transfer function of the network is obtained solving Eq. (3)-(5) which, in the limit in which the inputs to inhibitory neurons are much larger than threshold, is approximated by

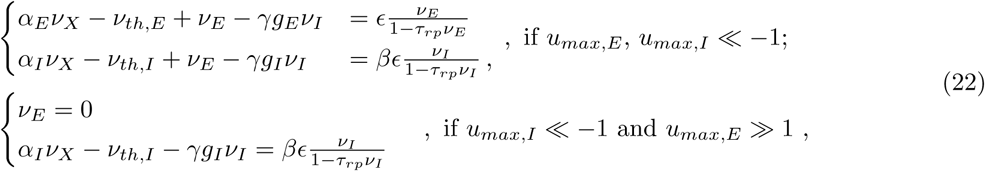

where 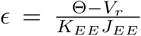 is the fraction of recurrent excitatory inputs that are needed to fire simultaneously to drive the membrane of an excitatory neuron from reset to threshold, 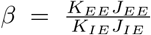 is a parameter measuring the ratio of total recurrent excitatory synaptic strength onto excitatory and inhibitory neurons, 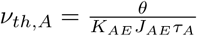 is the external firing rate needed to bring population *A* at firing threshold, in the absence of recurrent inputs.

Admissible solutions of Eq. (22), i.e. with rates in the range [0, 1*/τ*_*rp*_], provide a good approximation of the network transfer function. The advantage of using Eq. (22) is that it is polynomial in the rates and can be easily solved with an *ϵ* expansion of the form

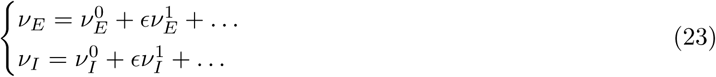

Moreover, as discussed above, 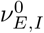 solutions describe, up to corrections of order 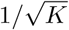, the network response for |*u*_*max*_| ∼ 1.

We now investigate the structure of the network transfer function using the *ϵ* expansion. To simplify expressions, in what follow we omit contributions coming from *ν*_*th,A*_. For *ϵ* = 0, there are two solutions of Eq. (22) in which both populations are not saturated. The first of these solutions, which will be called s1 or regular, has the first two terms of the expansion given by

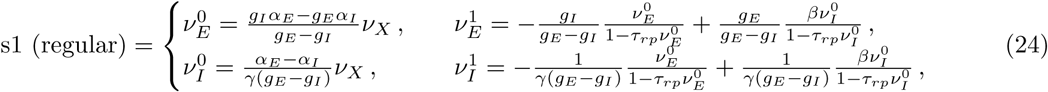

This solution corresponds to the classic ‘balanced’ solution [1; 2]. In the strong coupling limit, regular solutions feature excitatory and inhibitory rate increasing linearly with input strength. For finite coupling, the network transfer function deviates from this linear scaling. In particular, the response of one population is supralinear (sublinear) if the first order correction is positive (negative). For instance, the excitatory rate is a supralinear function of inputs if

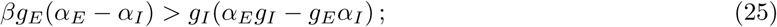

sublinear scaling appears if the above inequality is not satisfied. Analogous conditions holds for the inhibitory population. In agreement with this result, we find numerically that solutions of Eq. (3)-(5) which approach s1 solutions for *ϵ* = 0 transition from sublinear to supralinear as *β* increases (Fig. 4A).

The second unsaturated solution of Eq. (22) obtained for *ϵ* = 0 will be called s2 or supersaturated and is given by

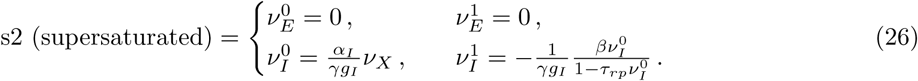

In the strong coupling limit, the inhibitory rate increases linearly with the external drive while the excitatory population remains silent due to overwhelming inhibition. Applying the same approach as the one used for s1 solutions, we find that s2 solutions at finite coupling admit only sublinear scaling (Fig. 4B). Supersaturating solutions have been recently proposed to underlie nonlinear summation in cortex [23; 24]. We will discuss them in more detail in the following sections.

Up to now, we have assumed that only one of the two solution types appears for any given value of *ν*_*X*_. In what follows we discuss under what conditions the assumption is valid and what happens if these conditions are not met. In this analysis, a key role is played by saturated solutions of Eq. (22) which are of three types

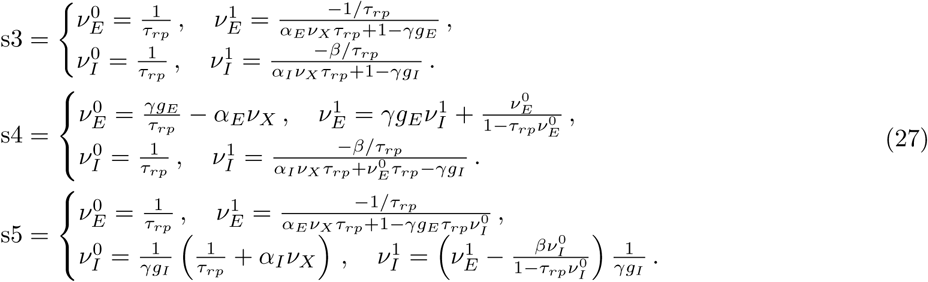

In the strong coupling limit, in s3 solutions, both populations are saturated; in s4 and s5 solutions, only one population is saturated while the other changes linearly with inputs.

To analyze solution admissibility, for any given value of *ν*_*X*_, we compute the rates predicted by the *E* expansion up to first order and investigate for what parameters each solution is within the [0, 1*/τ*_*rp*_] range. Conditions obtained for general values of *ν*_*X*_ are given in SI.. For simplicity, here we analyze results obtained for *ν*_*X*_ = 0; because of response continuity, these results are valid also in a range of sufficiently small external inputs. In this range of inputs, we find that there are seven possible scenarios, 3 with a single admissible solution (s1, s2 and s3), and 4 with three admissible solutions ({s1, s2, s3}, {s1, s2, s5}, {s1, s3, s4}, {s2-s3-s4}). When there are three solutions, we expect at least one of these solutions to be unstable; we will come back to this point during the analysis of network simulations and in the discussion.

We find that s1 solutions are the only one admissible if

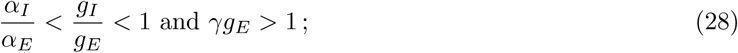

analogously, s2 solutions are the only one that appear if

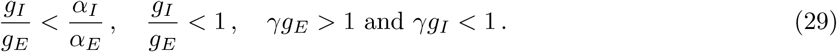

Note that the conditions in Eq. (29) are analogous to those for supersaturating solutions in the SSN [24]. The first two inequalities ensure suppression of excitatory rate with increasing input strength; reabsorbing *J*_*EA*_ into *α*_*A*_, they correspond to the conditions Ω_*E*_ < 0 and det *J* > 0 found in [24]. The last two inequalities prevent the existence of saturated solutions. Because of the presence of the refractory period in our model, the mathematical expressions are different in the SSN - the analogous condition in [24] being 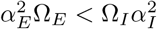. Any time Eqs. (28) or (29) are violated, multiple solutions appear. All the above-mentioned combinations of coexisting multiple solutions for *ν*_*X*_ ∼ 0 are shown explicitly in Fig. 4C-F. Note that the admissibility conditions derived in SI depends on the value of *ν*_*X*_. Therefore, as *ν*_*X*_ increases, the number and the identity of solutions is expected to change; examples of this phenomena are given in Fig. 4, where s1-s5 (Fig. 4D) or s1-s4 (Fig. 4E) solutions merge for *ν*_*X*_*τ*_*rp*_ ∼ 0.05 leaving only one admissible solution for larger inputs. Using the relations derived in SI, we also derive relations between solutions that must be satisfied for arbitrary values of *ν*_*X*_. First, we find that solutions s3 and s5 are mutually exclusive and can never appear at the same time. Second, any time that solution s1 and s2 are admissible at the same time, s4 solutions are not admissible. These two results imply that, for any value of *ν*_*X*_ at most three saturation-generated solutions can coexist. Finally, for large *ν*_*X*_, only s3 solutions are admissible, i.e. as the input intensity increases, both populations eventually saturate.

To summarize, we have shown that in model B at finite coupling, the network response can have one or multiple solutions. We have provided a general framework which, given the network parameters, predicts the expected number of solutions and their nonlinearities at finite coupling.

### Response-onset nonlinearities

In this Section, we characterize nonlinearities generated at response onset, first in model A and then in model B. In both models we show that, at response onset, one or multiple firing rates can coexist while, at larger input stimuli, response approaches the balanced-solutions described in the previous section.

#### Model A

To understand the possible nonlinearities appearing at response onset in model A, we rely on Eq. (15), whose solution provide a good approximation of the network response for *u*_*max*_ ≫ 1. In this regime, we were not able to find a useful perturbative expansion and hence we will use a different approach, analyzing how the response evolves as a function of input strength *ν*_*X*_.

For small enough *ν*_*X*_, effects of recurrent interactions in the network are negligible, feedforward inputs dominate over recurrent inputs, and the transfer function is given by

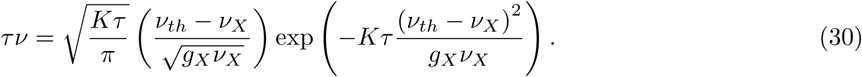

As shown in Fig. 5A, Eq. (30) captures the response for small rates: response rises exponentially as *ν*_*X*_ approaches *ν*_*th*_.

As *ν*_*X*_ increases, the network rate increases and recurrent inputs become relevant. In particular, recurrent inputs impacts network response as soon as their contributions to current mean or noise are of the same order as feedforward input. Specifically, using Eq. (15) we find that this happens as soon as *ν* approaches the minimum between (*ν*_*X*_ − *ν*_*th*_)/(1 − *gγ*) and *ν*_*X*_*g*_*X*_/(1 + *g*^2^*γ*). At this point, the transfer function is found solving the implicit Eq. ((3)-(5)) (or its simplified form Eq. (15)). We find numerically (see Fig. 5B-D) that, up to the point at which the membrane potential is close to threshold (*u*_*max*_ ∼ 1), there is either a single solution, or three solutions to the equations for the population firing rate. When a unique solution is present, firing increases supralinearly with inputs. For larger input values, the transfer function is determined by the linear response of the balanced-state regime.

As pointed out in [30; 31], multiple solutions emerge because of the positive feedback induced by the variance of the fluctuations in recurrent synaptic inputs and can be understood as follows. Let’s consider the fictitious dynamics given by [31]

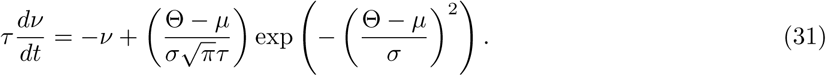

The fixed points of the above equation gives the network response at fixed input, i.e. the network transfer function; when multiple solutions are present, one solution must be unstable. This means there must be a solution for which the linearized dynamics

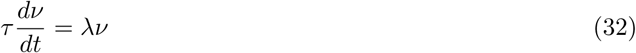

has *λ* > 0, with

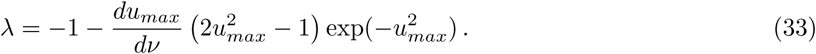

Since the above equation is valid for *u*_*max*_ ≫ 1, multiple solutions can be generated only if the derivative of *u*_*max*_ with respect to *ν* is sufficiently large and negative. This derivative is given by

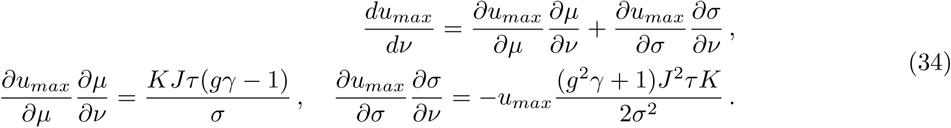

The above equation shows that, in an inhibition dominated network (*gγ* > 1), recurrent input mean and fluctuation have opposite effects; while the mean inputs provide negative feedback and hence cannot generate more than one solution, fluctuations provide positive feedback and therefore can potentially lead to multiple solutions.

This is verified in Fig. 5A, where we compare network response computed from Eq. ((3)-(5)) with and without either recurrent input mean (*μ*_*rc*_ = *ν*(1 − *gγ*)) or noise 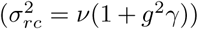); it show that noise is the key factor to produce multiple solutions.

We characterize the number of solutions numerically, since the nonlinearities make an analytic approach unfeasible. First we note that, out of the 5 model parameters in Eq. (15) (*τ, K, g*_*X*_, *J*, and *g*) only 4 are relevant in determining the response structure, since *τ* fixes the overall scale of the rates. Results of the numerical investigation are shown in Fig. 5B-E. For all parameters, the low rate response rises exponentially up to the point at which recurrent inputs are no longer negligible. After this point, the network shows either supralinear increases or multiple solutions. Increasing *J* and *K* (Fig. 5B-C) decreases the response onset point, as expected by the definition of *ν*_*th*_, and lead to multiple solutions. Changing *g* (Fig. 5D) has little effect on response onset but it affects the slope of the balanced-solution. Finally, increasing *g*_*X*_ (Fig. 5E) reduces the likelihood of having multiple solutions; this is due to the fact that recurrent input noise becomes negligible compared to the feedforward input noise.

#### Model B

In this section we analyze the behavior of model B at response onset. As discussed above, at low input rates, responses are determined by feedforward inputs, and the contribution of recurrent interactions are negligible. At larger input rates, when at least one of the two population has *u*_*max*_ of order one, recurrent interactions are relevant, the network response is approximately linear and approaches one of the solutions derived in the previous section (e.g., when only one solution is present, regular or supersaturated). The response region connecting these two regimes is expected to be nonlinear; characterizing possible nonlinearities is the goal of this section. Unlike the case of model A, however, the large number of parameters makes an extensive exploration of possible behaviors unfeasible. Therefore, we focus our investigation on the role of coupling strength, which in model A was found to have a major role in determining the type of response nonlinearity.

Examples of excitatory and inhibitory activity for regular and supersaturated solutions are shown in Fig. 6. We find that, in the region of inputs connecting response onset to the balanced solution, rates can either have a unique or multiple solutions. When only a unique solutions is present, response increases supralinearly in the regular case (Fig. 6A) and has a supralinear increase which eventually becomes sublinear in the supersaturated case (Fig. 6B). In model B, there are two independent ways in which coupling strength can be modified to affect these responses: uniform change (e.g. increase *K*_*EE*_ with *K*_*EE*_ = *K*_*IE*_) and relative change (e.g. fix *K*_*EE*_ and change *K*_*IE*_).

In regular solutions, a uniform increase in coupling strength reduces the region of nonlinear response, promoting a more sudden transition to linear scaling (Fig. 6A, first vs second columns). The same modification in supersaturated solutions reduces the peak response of excitatory cells and increases the likelihood of having multiple solutions (Fig. 6A, first vs second columns).

The main effect of changing the relative coupling is to move the relative onset point of the two populations, since it controls the ratio of *ν*_*th,E*_ and 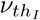. For instance, increasing *K*_*EE*_ at fixed *K*_*IE*_ makes the excitatory population respond at lower input rates than the inhibitory population. For small modifications, this leads to a larger region of supralinear response (e.g. first column of Fig. 6B light blue vs yellow lines). For larger modifications, it produces multiple solutions, since the excitatory population is unstable on its own (e.g. first column of Fig. 6B red line). Finally, we note that similar effects could be produced modifying the relative onset point of the two populations through changes in other parameters, such as *J*_*EE*_ of *J*_*IE*_.

In model B, there are two independent sources of positive feedback which can generate multiple solutions: noise and excitatory-to-excitatory connectivity. Which of these sources generates the observed multiple responses? We already know that, when network structure reduces model B to model A, i.e. when excitatory and inhibitory populations have the same properties, multiple solutions are generated by noise feedback. Are noise-generated multiple solutions limited to this specific case or a more general feature of model B? To answer this question, inspired by model A, we computed numerically the network response with and without recurrent noise, for different values of network parameters; results are shown in Fig. 7. At all coupling values, results along the line *g*_*E*_ = *g*_*I*_ are analogous to model A, as expected. At strong coupling, for a significant fraction of parameter space, the presence of multiple solutions is generated by noise, as they disappear when noise is removed. Interestingly, at weak coupling, noise seems to stabilize activity in a finite fraction of the parameter space; this phenomenon could be due to the larger increase in inhibitory response produce by its positive feedback coming from recurrent noise of inhibitory population.

These results show that, in model B, response onset is nonlinear and that a full characterization of response nonlinearities requires recurrent noise to be included in the model. This represents a qualitative difference with respect to rate models, where recurrent noise is not taken into account. In the next section we show how the results obtained so far compare with rate models.

### Comparison with network simulations and rate models

Results on response nonlinearities described up to this point have been obtained using a mean field analysis of networks of spiking neurons. In this section, we compare these results to other two approaches widely used in the study of neural networks: network simulations and rate models.

#### Network simulations

We perform numerical simulation in networks with either uniform or Erdös–Rényi connectivity. In the uniform case, all neurons receive exactly the same numbers of external, recurrent excitatory and recurrent inhibitory connections, i.e. there is a fixed in-degree for all types of connections. In the Erdös-Rényi (ER) case, the adjacency matrix specifying the existence of a synapse from any presynaptic to any (distinct) post-synaptic neuron is composed of i.i.d. Bernoulli variables, with connection probability given by ratio between mean number of connections and number of neurons in the presynaptic population. This leads to fluctuations in numbers of in-degrees between neurons which generates, in the balanced limit, a wide distribution of firing rates [1; 2; 3; 5]. The mean-field theory we have used so far assumes a fixed in-degree. Simulations of networks with uniform connectivity are useful to check that MF results are accurate, in spite of all the involved approximations (see Materials and Methods); simulations of ER networks allow us to assess the robustness of our results to heterogeneities.

Network simulations were performed with the simulator BRIAN2 [32] using networks of *N*_*E*_ = 11 *K*_*EE*_ excitatory and *N*_*I*_ = 11 *K*_*EI*_ inhibitory neurons, receiving excitatory inputs from ensembles of *N*_*EX,IX*_ = 11 *K*_*EX,IX*_ independent Poisson units firing at rates *ν*_*EX,EI*_, respectively. We used uniformly distributed delays of excitatory and inhibitory synapses. Delays were drawn randomly and independently at each existing synapse from uniform distributions in the range [0, 100]ms (E synapses) and [0, 1]ms (I synapses) [31]. These extreme values ensure no synchronized oscillations are present (see Discussion). For fixed network parameters, rates were computed starting from a given initial condition and gradually increasing the external drive. We explored different initial conditions to check for multiple solutions. Stationary responses obtained for different representative parameter sets are shown in Fig. 8. For connectivity parameters leading to saturation nonlinearity, results of the network simulations closely match predictions of the mean field theory. In the case in which multiple solutions are expected from the mean field theory, either generated at saturation or at response-onset, simulations capture the upper and lower branches of the response; the absence of the middle branch suggests that it corresponds to unstable fixed points of the dynamics. Simulations obtained for parameters generating supersaturation in the mean field model are found to depend on the connectivity structure. For uniform connectivity, simulation follows the same trend of the mean field prediction, with inhibitory rate increasing monotonically while excitatory rate show an increase with inputs at low intensity and suppression at larger inputs. We note that, although this mean response follows the trend of the mean field prediction, the network showed oscillatory activity in the region of maximum excitatory response despite the broad distribution of synaptic delays. In the case of random connectivity, on the other hand, we found that the suppression of excitatory response is not present. This lack of supersaturation at the population level is generated by an heterogeneity in responses of excitatory cells, with 66% of cells which are silent while the remaining 33% show a weak increase in rate with the external drive. Despite this difference, a common features of supersaturating solutions in networks of spiking neurons, observed both in the mean filed theory and in simulations, is that the peak response of excitatory cells is small. The intuitive reason why firing rates of excitatory neurons are generically small in this scenario is that rates go to zero in the strong coupling limit.

These numerical results validate the analysis derived above as a good description of the response in spiking networks, with the only exception given by the lack of supersaturation in networks with random connectivity. In what follows we compare how rate models compare to results obtained in spiking networks.

#### Rate models

Rate models are characterized by a fixed relation between input current *μ* and firing rate response (here referred to as *f* − *μ* curve); this is not the case in spiking networks, where the firing rate also depends on the noise level which, in turns, depends on the input rate. Here we focus on two specific models: a ‘Ricciardi’ model, in which the single unit transfer function is given by the Ricciardi nonlinearity at fixed noise value, and the SSN [24], where the single unit transfer function is a power law with an exponent that is larger than one; a mathematical definition of these models is given in the Methods section. The comparison between different models is performed computing responses in networks of given input structure *α*_*E,I*_ and of recurrent inhibition *g*_*E,I*_; this choice ensures the same balanced-solutions in all models and limits differences to the nonlinear response regions. In the Ricciardi-model, parameters are as in spiking network, the only exception is the noise level, which must be specified. We fix this to be approximately the value observed in the spiking-network model at threshold. In the SSN, we fix parameters as in [24], i.e. we assume a powerlaw nonlinearity with exponent *n* = 2 and proportionality constant *k* = 0.04.

For networks producing saturation nonlinearities (Fig. 8 first and second rows), the Ricciardi-model recapitulates quantitatively responses observed in spiking networks. As mentioned during our analysis of saturation nonlinearities, this is due to the fact that in spiking network the response in the balanced-regime and for larger rates depends weakly on the noise level. It follows that the agreement between spiking networks and the Ricciardi model is valid for all saturation nonlinearities discussed in model A and B. The SSN, on the the hand, shows substantial discrepancies. In particular, for network structure generating sublinear response in spiking network (Fig. 8 first row), the SSN responds linearly; in the parameter conditions in which spiking network shows multiple solutions at zero inputs and maximum firing at large inputs (Fig. 8 second row), the SSN shows supersaturation. These discrepancies are expected, since the SSN does not include a refractory period, i.e. the ingredient needed to generate saturation nonlinearity. While these nonlinearities can be captured by adding a refractory period to the model, our results show that consequences of the refractory period appear at rates that are much lower than 1*/τ*_*rp*_. First, for moderate coupling, response is nonlinear for rates that are far from the maximum firing rate. Second, saturation nonlinearities limit the parameter space in which the network has a unique solution. In particular, the network structure used in Fig. 8 second row is the one used in [24; 25], where a large region of nonlinear responses has been used to explain nonlinearities observed in input summations in the cortex. Our results show that these parameters produce multiple solutions (specifically having *g*_*E*_ < 1*/γ* generates the coexistence of s2, s3 and s4) and saturation when the refractory period is taken into account. This saturation of the two population in s2 solutions comes from the fact that, at large inputs, the inhibitory population saturates while the external input to excitatory cells (*α*_*E*_*ν*_*X*_) keeps increasing. As showed during the analysis of saturation nonlinearities in model B, supersaturation without additional solutions can be generated imposing *g*_*E*_ > 1*/γ*, i.e. with larger recurrent inhibition to excitatory cells. Note that, at a given value of inhibitory rates, this constraint limits the peak amplitude that the excitatory population can reach in supersaturating solutions.

We next focus on response-onset nonlinearities. In spiking networks, rates at responses onset can either increase supralinearly or go through a region with multiple solutions; for larger inputs, rates approach the balanced solutions. Example of these transitions are shown in the last three rows of Fig. 8. For network structure leading to supersaturating solution in spiking networks, the same qualitative behavior is observed in rate models but quantitative differences are observed. First, because of feedback coming from recurrent noise, the supralinear region at response onset is stiffer in spiking network. Indeed, the response in the Ricciardi-model, where the noise amplitude is fixed and there is no positive feedback coming from noise, is much less steep. A second major quantitative difference is that the peak excitatory rate is smaller in spiking network and Ricciardi model with respect to the SSN; we will come back to this point at the end of this section. For network structures leading to multiple solutions at response-onset, we should distinguish two cases: mean and noise generated multiple solutions. For some parameters, positive feedback due to mean recurrent input generates multiple solutions; this mechanism is also present in the Ricciardi model, and also in the SSN (Fig. 8, fourth row). On the other hand, when multiple solutions are generated by noise feedback in spiking networks, there are no multiple solutions in rate models (Fig. 8, fifth row); this is expected because in rate models there is no positive feedback generated by recurrent noise.

#### Effects of coupling strength in the supersaturating scenario

We have shown that, in spiking networks, supersaturating solutions tend to have a lower peak response than in the SSN (Fig. 8, third row). Here we investigate how this result changes as a function of coupling strength We perform our investigation using the Ricciardi model, which can be more easily compared with the SSN, since they both have a fixed *f* − *μ* curve; similar results were found in the LIF network (data not shown).

The SSN [24] is characterized by a supralinear *f* − *μ* curve (specifically, a power-law), which generates an effective coupling strength between neurons that increases monotonically with rates. Writing the stationary condition for the two populations (*E, I*) as

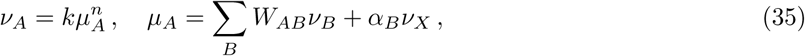

the effective weight from population *B* to population *A* is given by

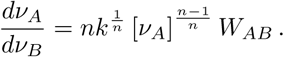

Therefore, as *ν*_*X*_ increases, no matter how small the *W* s are, there is always be a value at which the effective coupling becomes strong enough to trigger an instability of the excitatory network, which is eventually stabilized by inhibition. As shown in [24], the excitatory response is suppressed at larger *ν*_*X*_, i.e. supersaturation appears, if the connectivity connectivity is such that

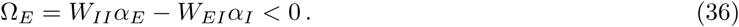

In the Ricciardi model, the *f* − *μ* features an expansive nonlinearity, with an exponent larger than two, at low *μ*, and becomes linear as *μ* increases (Fig. 9B). Applying Eq. (35) locally, around a given value of *μ*, shows that effective coupling strength increases with activity only up to the linear region of the *f* − *μ* curves, and saturates in that linear region. Note that in this figure we use *τ*_*rp*_ = 0. In the presence of a refractory period *τ*_*rp*_ > 0, the effective coupling strength reaches a maximum and then gradually decreases as the neuron gets closer to saturation. Therefore, in the Ricciardi model, the excitatory instability can be generated only if the effective weights become large enough before the point at which the *f* − *μ* curve becomes linear. On the other hand, if the effective weights are not large enough to destabilize excitation when the *f* − *μ* curve enters the linear region, excitatory instability, and hence supersaturation, will not appear even for larger current values. In agreement with this argument we find that, while a moderate reduction in coupling strength increases the peak excitatory rate, supersaturation eventually disappears as coupling strength decreases (Fig. 9A). At the same time, the instability point of the excitatory subnetwork, defined as the point at which 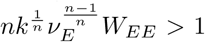, approaches the linear part of the *f* − *μ* (Fig. 9B). Interestingly, supersaturation disappears for strengths which still generate excitatory instability, consistent with the fact that instability is a necessary but not sufficient condition for supersaturation [24].

**FIG. 9.**
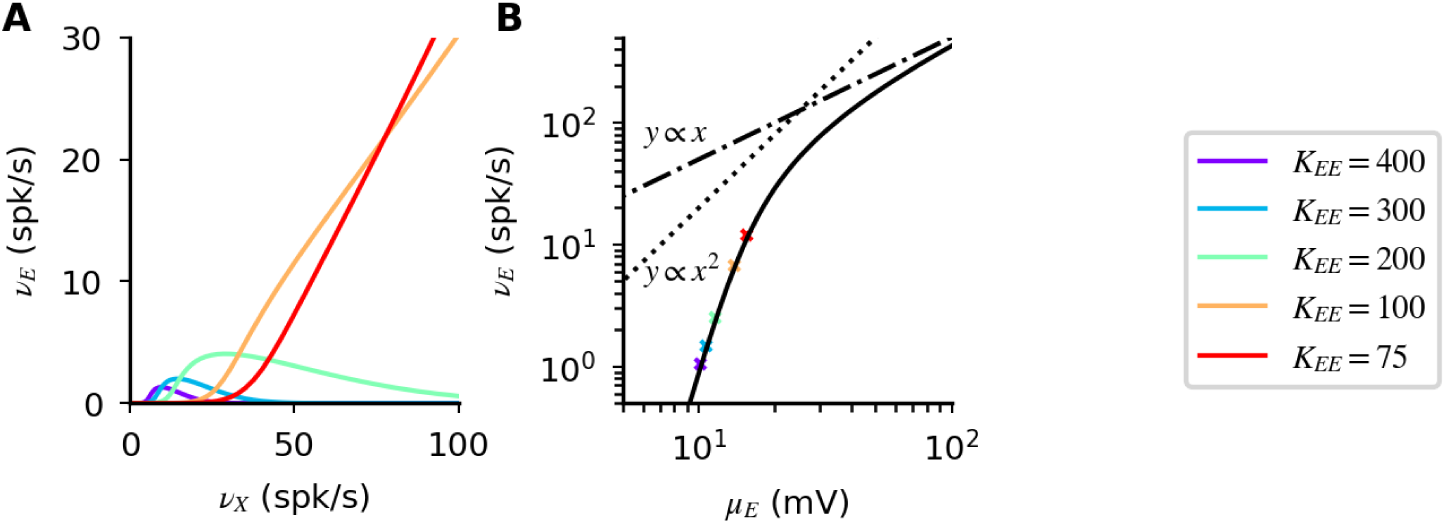
Disappearance of supersaturation as the coupling strength decreases. (**A**) Excitatory response in Ricciardi model for different coupling strength. (**B**) *f* − *μ* curve of Ricciardi model (black line); it starts supralinearly and becomes linear for large currents. Linear (dash-dotted line) and quadratic (dotted line) scalings are shown as references. Crosses show the location at which the excitatory subnetwork becomes unstable if inhibition is kept constant. Simulation parameters: *α*_*E*_ = *α*_*I*_ = 1; *g*_*E*_=4.;*g*_*I*_=2.7; *J*_*EE*_ = *J*_*IE*_ = 0.2mV. To simplify comparison with SSN, we assumed *τ*_*rp*_ = 0.

To summarize, in the Ricciardi model, the peak excitatory response of supersaturating solutions increases for a moderate decrease in coupling strength; a more significant reduction leads to the disappearance of supersaturation.

## DISCUSSION

In this work, we have investigated nonlinearities emerging in networks of spiking neurons at finite coupling, showing that these are of two types: response-onset and saturation. The network response transitions between these two nonlinearities as feedforward input increases; for intermediate inputs, the response matches that of the balanced-state model up to corrections of order 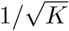. Importantly, the influence of saturation emerges already at rates that are much lower than the single neuron maximum response, producing sublinear or supralinear response and affecting the number of solutions at low inputs. Therefore, both types of nonlinearities can be relevant at activity levels observed in the brain. Our results have been obtained using a mean-field analysis, but are also confirmed in numerical simulations of large networks. Finally, we have analyzed which of the features of the response of spiking networks can be recapitulated by rate models.

Nonlinear operations are thought to underlie contrast dependent input summations observed in cortex, in surround suppression [17; 18] and normalization [19; 20]. These phenomena have been recently explained by the SSN model, using parameters of network connectivity that lead to supersaturating responses [25]. Supersaturating solutions show linear/supralinear response at low inputs and sublinear response at intermediate inputs; i.e. nonlinearities analogous to those observed in cortex for low and high inputs, respectively [17; 18; 19; 20]. In this work, we have shown that all the nonlinearities generated in the SSN are present in spiking networks, but substantial differences exist between the two models. In particular, in spiking networks, supersaturating solutions with nonzero excitatory rates exist only for intermediate coupling strength and exhibit much smaller firing rates than in the SSN. This result seems to question the applicability of supersaturation to explain some nonlinearities observed in the cortex, since they emerge over rate changes up to tens of *spk/s* [19]. Note however that we cannot exclude having missed small regions in parameter space for which more robust supersaturation could occur. Also, one cannot exclude that this issue might be alleviated by the use of more detailed spiking neuron models. Moreover, as showed in Fig. 9, although there is no supersaturation in spiking networks at weak coupling, there still can be linear/supralinear summation at low inputs and sublinear summation at larger inputs, i.e. the key ingredients used by the SSN to explain cortex phenomenology. Finally, the presence or absence of supersaturation discussed here refers to cases in which the strength of a single stimulus is varied. However, nonlinearities modeled by the SSN primarily involve summation of responses to two *different* stimuli (e.g. two stimuli in the receptive field or stimuli in center and surround). All of these nonlinear phenomena arise in the SSN when structured connectivity is considered, both in regimes featuring non-supersaturating and supersaturating responses [24; 25; 33] (Ken Miller, private communication). This suggests that structured connectivity can yield nonlinear behaviors not captured by the analysis discussed here and which need further investigation.

Our analysis suggests a different explanation of nonlinear summation in cortex. We found that, for finite coupling, saturation nonlinearities generate an activity dependent summation: at low intensity, inputs are summed linearly; summation becomes increasingly sublinear as the intensity increases. This nonlinearity emerges at relatively low rates and evolves gradually over activity levels which span few hundreds *spk/s*, a feature which compares favorably with some experimental results [19]. Moreover, unlike the SSN, this model does not require supralinear input summation at low activity level, a feature which has (to our knowledge) not been widely reported so far. Further experimental studies are needed to understand which mechanism underlies nonlinear input summation in cortex.

We have found that, in networks of spiking neurons, firing irregularity, i.e. the CV of the ISI, decreases monotonically in regular solutions for sufficiently large inputs. This effect is consistent with the stimulus-driven suppression of variability that has been reported in various experiments [34; 35; 36; 37; 38]. In our model, regardless of the specific parameters used, variability suppression emerges as the mean input approaches firing threshold, a robust prediction which is consistent with results from intracellular recordings [34; 35; 37; 39]. This mechanism differs qualitatively from alternative scenarios that have been proposed [33; 40]. Additional experimental and theoretical investigations are needed to understand which of these possibilities underlies the stimulus-driven suppression of variability observed in the brain.

Our analysis revealed that multiple solutions can appear at response-onset, produced by noise-driven or mean-driven positive feedback, and at larger rate values, generated by saturation of one population. In both cases, neural response can assume one of three possible values; combining these mechanisms, we find that, in spiking networks, there can be up to five coexisting solutions for every fixed value of the external drive. Even though the stability of these solutions has not been investigated here, network simulations suggests that in the case three solutions coexist, up to two of them can be stable (the one with lowest and highest rates), while the intermediate solution is unstable. In the case of 5 coexisting solutions, up to three of them can be stable (the ones with lowest, 3rd highest and highest rates) while intermediate solutions are again unstable. A recent work [41] analyzed the number of coexisting solutions in the SSN, showing that the model can support up to four fixed points at fixed external drive, with only two being stable. The larger number of solutions found in spiking networks is generated by single neuron saturation.

Multiple studies have found that fixed points can also become unstable due to oscillatory instabilities, depending for instance on the distribution of synaptic delays [4; 42], and these oscillations can destroy bistability [4; 31]. For instance, it has been shown that, for certain parameters, the asynchronous state is stable only for intermediate values of the external drive, while synchronous states exist both for low external drive [4; 31], and high external drive [4; 43; 44]. This pattern bears similarities with experimental data obtained in primary visual cortex, both in monkeys [39] and mice [45]. In addition, synchronous oscillations at low inputs can generate an effective supralinear transfer function in a region with multiple (unstable) fixed points coexist, through time average of network activity. In this context, our classification of multiple solutions could be used as a starting point to systematically classify the possible oscillatory regions that can emerge in a network.

We developed a new approach to study network response at finite coupling strength, allowing us to classify all possible deviations from linear response. Although this framework has been derived in the case of one excitatory and one inhibitory population, it can be readily generalized to study models with multiple interacting populations. An important application concerns the case of networks with one excitatory and multiple inhibitory populations, e.g. SOM, PV and VIP cells. The interaction between these neuron types has been recently implicated, among other phenomena, in the top-down modulation of activity during locomotion [46; 47; 48; 49; 50]. In particular, the observed change of modulation with context (dark vs visual stimuli) has been explained with a rate model with supralinear *f* − *μ* curve, assuming a change in baseline activity (low vs high) [51]. As we have shown in the present work, other scenarios exist close to response onset in networks of spiking neurons, which might lead to alternative explanations of the observed phenomena. Understanding the role of nonlinearities in this and other phenomena will require additional theoretical and experimental work.

Nonlinearities studied in this contribution are generated uniquely from the interplay between single neuron properties and network connectivity, using synapses with fixed efficacy. Synapses in the brain exhibit changes at multiple time scales, which could affect qualitatively the picture derived here. Further theoretical and experimental investigations are needed to understand how the interplay between these mechanisms shapes response properties of networks in the brain.

To conclude, we have shown that a simple network of interacting excitatory and inhibitory spiking neurons shows a rich repertoire of nonlinearities. Most neural systems receive information from multiple inputs. This is true not only in sensory systems, where most of the currently available data come from, but also in associative areas, where information coming from different sensory systems is combined with an animal cognitive state to make decisions and drive behavior. The general rules with which different inputs are combined for these and other computations are still the subject of debate; our work provides the basis to systematically study the biophysical constraints on these operations.

## ACKNOWLEDGMENTS

Supported by the NIMH Intramural Research Program and by NIH BRAIN U01 NS108683. We thank K. Miller for discussions and for his comments on the manuscript. This work used the computational resources of the NIH HPC Biowulf cluster (http://hpc.nih.gov).

## SUPPORTING INFORMATION

### Admissibility conditions for model B solutions at finite coupling

In the main text, we have shown that there are 5 solutions which are admissible in the strong coupling limit of model B. At finite coupling, only solutions producing rates below saturation are admissible. Using this constraint, for every solution described in Eqs (24), (26) and (27), we derive the following admissibility conditions.

Solutions of s1 type appear if connectivity is such that

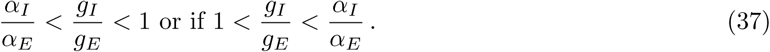

Moreover, the external input must satisfy the condition

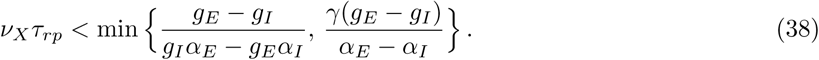

i.e. none of the two populations is saturated.

Solutions of s2 type appear only if

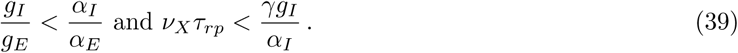

Solutions of s3 type appear only if

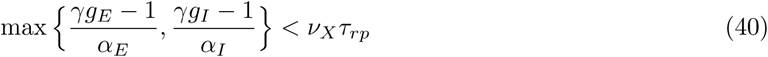

Solutions of s4 type appear only if

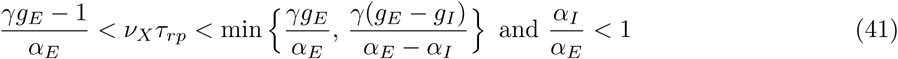

or if

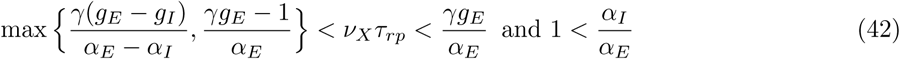

Solutions of s5 type appear only if

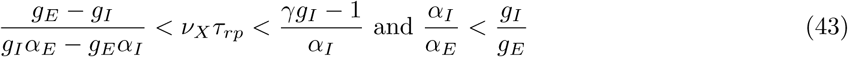

or if

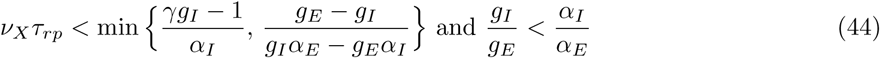

Combining the above inequalities we find the following relations between the different solutions

- Conditions on s3 and s5 solutions are mutually exclusive, which implies that only one of the two solutions type can appear at any given value of *ν*_*X*_.
- When solutions s1 and s2 coexists, s4 solutions are excluded.

Combining these two results we find that for any value of *ν*_*X*_ there can be at most three acceptable solutions.

## REFERENCES

[1] C van Vreeswijk and H Sompolinsky. Chaos in neuronal networks with balanced excitatory and inhibitory activity. Science, 274(5293):1724–6, 1996.

[2] C van Vreeswijk and H Sompolinsky. Chaotic balanced state in a model of cortical circuits. Neural computation, 10(6):1321–71, 1998.

[3] D J Amit and N Brunel. Model of global spontaneous activity and local structured activity during delay periods in the cerebral cortex. Cerebral cortex (New York, N.Y. : 1991), 7(3):237–252, 1997.

[4] N Brunel. Dynamics of sparsely connected networls of excitatory and inhibitory neurons. Journal of Computational Neuroscience, 8:183–208, 2000.

[5] Alex Roxin, Nicolas Brunel, David Hansel, Gianluigi Mongillo, and Carl van Vreeswijk. On the distribution of firing rates in networks of cortical neurons. Journal of Neuroscience, 31(45):16217–16226, 2011.

[6] A. Renart, J. de la Rocha, P. Bartho, L. Hollender, N. Parga, A. Reyes, and K. D. Harris. The Asynchronous State in Cortical Circuits. Science, 327(5965):587–590, jan 2010.

[7] Robert Rosenbaum, Matthew A Smith, Adam Kohn, Jonathan E Rubin, and Brent Doiron. The spatial structure of correlated neuronal variability. Nature Neuroscience, 20:107 EP –, 10 2016.

[8] Ran Darshan, Carl van Vreeswijk, and David Hansel. Strength of correlations in strongly recurrent neuronal networks. Phys. Rev. X, 8:031072, Sep 2018.

[9] Cody Baker, Christopher Ebsch, Ilan Lampl, and Robert Rosenbaum. Correlated states in balanced neuronal networks. Phys. Rev. E, 99:052414, May 2019.

[10] R J Douglas and K A Martin. A functional microcircuit for cat visual cortex. The Journal of physiology, 440:735–769, 1991.

[11] WR Softky and C Koch. The highly irregular firing of cortical cells is inconsistent with temporal integration of random epsps. Journal of Neuroscience, 13(1):334–350, 1993.

[12] W Bair, C Koch, W Newsome, and K Britten. Power spectrum analysis of bursting cells in area mt in the behaving monkey. Journal of Neuroscience, 14(5):2870–2892, 1994.

[13] Tomáš Hromádka, Michael R DeWeese, and Anthony M Zador. Sparse representation of sounds in the unanesthetized auditory cortex. PLOS Biology, 6(1):1–14, 01 2008.

[14] Daniel H. O’Connor, Simon P. Peron, Daniel Huber, and Karel Svoboda. Neural activity in barrel cortex underlying vibrissa-based object localization in mice. Neuron, 67(6):1048 – 1061, 2010.

[15] Alexander S. Ecker, Philipp Berens, Georgios A. Keliris, Matthias Bethge, Nikos K. Logothetis, and Andreas S. Tolias. Decorrelated neuronal firing in cortical microcircuits. Science, 327(5965):584–587, 2010.

[16] Alessandro Sanzeni, Mark Histed, and Nicolas Brunel. Dynamics of networks of conductance-based neurons in the strong coupling limit. Computational and Systems Neuroscience (Cosyne), Mar 2018.

[17] F. Sengpiel, Arjune Sen, and C. Blakemore. Characteristics of surround inhibition in cat area 17. Experimental Brain Research, 116(2):216–228, Sep 1997.

[18] Uri Polat, Keiko Mizobe, Mark W Pettet, Takuji Kasamatsu, and Anthony M Norcia. Collinear stimuli regulate visual responses depending on cell’s contrast threshold. Nature, 391(6667):580–584, 1998.

[19] Hilary W. Heuer and Kenneth H. Britten. Contrast dependence of response normalization in area mt of the rhesus macaque. Journal of Neurophysiology, 88(6):3398–3408, 2002. PMID: 12466456.

[20] Tomokazu Ohshiro, Dora E Angelaki, and Gregory C DeAngelis. A normalization model of multisensory integration. Nature neuroscience, 14(6):775–782, jun 2011.

[21] Bruno A. Olshausen and David J. Field. How close are we to understanding v1? Neural Computation, 17(8):1665–1699, 2005.

[22] Gianluigi Mongillo, David Hansel, and Carl van Vreeswijk. Bistability and spatiotemporal irregularity in neuronal networks with nonlinear synaptic transmission. Phys. Rev. Lett., 108:158101, Apr 2012.

[23] Erez Persi, David Hansel, Lionel Nowak, Pascal Barone, and Carl van Vreeswijk. Power-law input-output transfer functions explain the contrast-response and tuning properties of neurons in visual cortex. PLOS Computational Biology, 7(2):1–21, 02 2011.

[24] Yashar Ahmadian, Daniel B Rubin, and Kenneth D Miller. Analysis of the stabilized supra-linear network. Neural Computation, 25(8):1994–2037, 2013.

[25] Daniel B. Rubin, Stephen D. VanHooser, and Kenneth D. Miller. The stabilized supralinear network: A unifying circuit motif underlying multi-input integration in sensory cortex. Neuron, 85(2):402–417, 2015.

[26] Alessandro Sanzeni, Mark Histed, and Nicolas Brunel. Emergence of nonlinear computations in spiking neural networks with linear synapses. Neuroscience Meeting Planner, Nov 2017.

[27] L. M. Ricciardi. Diffusion processes and related topics in biology. Springer-Verlag Inc, Berlin; New York, 1977.

[28] Nicolas Brunel, Vincent Hakim, and Magnus JE Richardson. Single neuron dynamics and computation. Current Opinion in Neurobiology, 25:149 – 155, 2014. Theoretical and computational neuroscience.

[29] Francesca Barbieri and Nicolas Brunel. Irregular persistent activity induced by synaptic excitatory feedback. Frontiers in Computational Neuroscience, 1:5, 2007.

[30] Alfonso Renart, Rubén Moreno-Bote, Xiao-Jing Wang, and Néstor Parga. Mean-Driven and Fluctuation-Driven Persistent Activity in Recurrent Networks. Neural Computation, 19:1–46, 2007.

[31] Elisa M. Tartaglia and Nicolas Brunel. Diversity of dynamical states in sparsely connected networks of lif neurons with strong inhibition. Preprint, 2017.

[32] Marcel Stimberg, Romain Brette, and Dan Fm Goodman. Brian 2, an intuitive and efficient neural simulator. eLife, 8:e47314, aug 2019.

[33] Guillaume Hennequin, Yashar Ahmadian, Daniel B. Rubin, Máté Lengyel, and Kenneth D. Miller. The dynamical regime of sensory cortex: Stable dynamics around a single stimulus-tuned attractor account for patterns of noise variability. Neuron, 98(4):846–860.e5, 2018.

[34] Ian M. Finn, Nicholas J. Priebe, and David Ferster. The emergence of contrast-invariant orientation tuning in simple cells of cat visual cortex. Neuron, 54(1):137–152, 2007.

[35] James F. A. Poulet and Carl C. H. Petersen. Internal brain state regulates membrane potential synchrony in barrel cortex of behaving mice. Nature, 454(7206):881–885, 2008.

[36] Mark M Churchland, Byron M Yu, John P Cunningham, Leo P Sugrue, Marlene R Cohen, Greg S Corrado, William T Newsome, Andrew M Clark, Paymon Hosseini, Benjamin B Scott, David C Bradley, Matthew a Smith, Adam Kohn, J Anthony Movshon, Katherine M Armstrong, Tirin Moore, Steve W Chang, Lawrence H Snyder, Stephen G Lisberger, Nicholas J Priebe, Ian M Finn, David Ferster, Stephen I Ryu, Gopal Santhanam, Maneesh Sahani, and Krishna V Shenoy. Stimulus onset quenches neural variability: a widespread cortical phenomenon. Nature neuroscience, 13(3):369–378, 2010.

[37] Luc J. Gentet, Michael Avermann, Ferenc Matyas, Jochen F. Staiger, and Carl C. H. Petersen. Membrane potential dynamics of gabaergic neurons in the barrel cortex of behaving mice. Neuron, 65(3):422–435, 2010.

[38] Mo Chen, Linyu Wei, and Yu Liu. Motor preparation attenuates neural variability and beta-band lfp in parietal cortex. Scientific Reports, 4:6809 EP –, 10 2014.

[39] Andrew Y. Y. Tan, Yuzhi Chen, Benjamin Scholl, Eyal Seidemann, and Nicholas J. Priebe. Sensory stimulation shifts visual cortex from synchronous to asynchronous states. Nature, 509:226 EP –, 03 2014.

[40] Kanaka Rajan, Laurence F Abbott, and Haim Sompolinsky. Stimulus-dependent suppression of intrinsic variability in recurrent neural networks. BMC Neuroscience, 11(Suppl 1):O17–O17, 07 2010.

[41] Nataliya Kraynyukova and Tatjana Tchumatchenko. Stabilized supralinear network can give rise to bistable, oscillatory, and persistent activity. Proceedings of the National Academy of Sciences, 115(13):3464–3469, 2018.

[42] Nicolas Brunel and Vincent Hakim. Fast global oscillations in networks of integrate-and-fire neurons with low firing rates. Neural Computation, 11(7):1621–1671, 1999.

[43] Alberto Mazzoni, Nicolas Brunel, and Stefano Panzeri. How gamma-band oscillatory activity participates in encoding of naturalistic stimuli in random networks of excitatory and inhibitory neurons. BMC Neuroscience, 9(1):P115, 2008.

[44] Francesca Barbieri, Alberto Mazzoni, Nikos K. Logothetis, Stefano Panzeri, and Nicolas Brunel. Stimulus dependence of local field potential spectra: Experiment versus theory. Journal of Neuroscience, 34(44):14589–14605, 2014.

[45] Matthew J. McGinley, Stephen V. David, and David A. McCormick. Cortical membrane potential signature of optimal states for sensory signal detection. Neuron, 87(1):179 – 192, 2015.

[46] Cristopher M. Niell and Michael P. Stryker. Modulation of visual responses by behavioral state in mouse visual cortex. Neuron, 65(4):472 – 479, 2010.

[47] Georg B. Keller, Tobias Bonhoeffer, and Mark Hübener. Sensorimotor mismatch signals in primary visual cortex of the behaving mouse. Neuron, 74(5):809 – 815, 2012.

[48] Yu Fu, Jason M. Tucciarone, J. Espinosa, Nengyin Sheng, Daniel P. Darcy, Roger A. Nicoll, Z.Josh Huang, and Michael P. Stryker. A cortical circuit for gain control by behavioral state. Cell, 156(6):1139 – 1152, 2014.

[49] Janelle Mp Pakan, Scott C Lowe, Evelyn Dylda, Sander W Keemink, Stephen P Currie, Christopher A Coutts, and Nathalie L Rochefort. Behavioral-state modulation of inhibition is context-dependent and cell type specific in mouse visual cortex. eLife, 5:e14985, aug 2016.

[50] Mario Dipoppa, Adam Ranson, Michael Krumin, Marius Pachitariu, Matteo Carandini, and Kenneth D. Harris. Vision and locomotion shape the interactions between neuron types in mouse visual cortex. Neuron, 98(3):602 – 615.e8, 2018.

[51] Luis Carlos Garcia del Molino, Guangyu Robert Yang, Jorge F Mejias, and Xiao-Jing Wang. Paradoxical response reversal of top-down modulation in cortical circuits with three interneuron types. eLife, 6:e29742, ec 2017.

